# Centimeter-sized Human Vitreous Organoids from Pluripotent Stem Cells

**DOI:** 10.64898/2026.11.19.767433

**Authors:** Xiao Zhang, Kangxin Jin, Xiao Lin, Fanying Zeng, Mingxia Du, Hai-Long He, Dongdong Zhao, Yue Zhang, Wen Wang, Ping Liu, Changhao Bi, Zi-Bing Jin

## Abstract

The vitreous, the largest intraocular tissue, provides a transparent light path and maintains the intraocular homeostasis. The reconstitution of the vitreous *in vitro* or *in vivo* remains an unsolved biomedical challenge. Here, we discovered a common feature, the “growth edge”, constituted primarily of fibroblasts and their secreted proteins in human and mouse vitreous development. By leveraging the transforming growth factor beta (TGFβ) signaling, we then generated centimeter-scale vitreous-like organoids (VOs) through stepwise differentiation from human pluripotent stem cells (hPSCs). The VOs exhibited striking similarity to human vitreous in terms of physical transparency, vitreoretinal interface, molecular composition, biological profiles and regenerative transplantability. Lines of evidence supported that fibroblasts are the key cell hub for vitreous development. Finally, VOs with a *transthyretin (TTR)* mutation recapitulated the vitreous phenotypes of human patients with vitreous amyloidosis. Together, vitreous organoid technology opens avenues for future studies of human vitreous development, disease modeling, and regenerative therapeutics for a broad spectrum of human vitreoretinal diseases.

## Introduction

The vitreous (also called vitreous body), the largest intraocular tissue in the human eye, is a transparent, highly hydrated (>98% water) extracellular matrix (ECM) located between the lens and retina^1–3^. The vitreous is heterogeneous and can be divided into three regions, the basal vitreous, the vitreous cortex, and the central vitreous. The concentration of the ECM varies between these regions and generally decreases from the periphery to the center^4,5^. The ECM plays a major role in maintaining the gel state of the vitreous^6,7^ and consists mainly of collagens (e.g., the type II,V/XI and IX), non-collagenous structural proteins (e.g., fibrillin, GAPDH, tubulin), and some other macromolecules (e.g., fibronectin, laminin, glycosaminoglycan (GAG))^4,8^.

The structural function of vitreous is to maintain the optical clarity and support the round shape of the eye, however, its physiological function is much underappreciated. As the vitreous occupies the biggest volume of the eye ball, the development of vitreous plays a crucial role in regulating the size of the globe^9^. In a broad spectrum of human eye diseases, both high axial myopia and microphthalmia are associated with the abnormal vitreous development^10–12^. Many inherited congenital disorders are also associated with pathological vitreous, including vitreous amyloidosis caused by the transthyretin (TTR) gene mutations^13,14^, vitreoretinopathies caused by the mutations in *COL2A1*, *COL11A1*, *etc.*^15,16^, leading to visual impairments and even blindness in severe cases. Moreover, since the vitreous is adjacent to the retina, vitreous abnormalities including liquefaction, hemorrhage and posterior vitreous detachment (PVD)^17–21^, are closely related to retinopathy (e.g. retinal detachment, macular hole, neovascularization and proliferative diabetic retinopathy), the leading cause of blindness worldwide.

Therefore, vitrectomy, i.e., removal of part of the vitreous, is commonly used in clinical practice for slowing the progression of vitreoretinopathies and preventing permanent vision loss^22,23^. However, the vitreous does not have the ability to regenerate. An artificial vitreous substitute is needed to maintain the ocular structure. An ideal vitreous substitute should possess the physical features (transparence) of the vitreous and achieve the biocompatibility and long-term intraocular permanence, which has not yet been realized^24–26^. The vitreous-like organoids (VOs), derived from human pluripotent stem cells (PSCs), including embryonic stem cells (ESCs) and induced pluripotent stem cells (iPSCs), would fulfill these requirements. In addition, VOs could serve as disease models to uncover the pathogenesis causing human vitreous or vitreoretinal disorders.

The cellular and molecular mechanisms regulating mammalian vitreous development *in vivo* are still unclear, particularly in humans. For example, how the vitreous gel state is formed, maintained, and later remodeled remains unresovled^4,8^. Fibroblasts, hyalocytes, lens epithelial cells, retinal Müller glial cells and pigment cells, and ciliary body cells have been suggested to contribute to vitreous formation, however, the cellular origin of vitreous macromolecules remains a mystery^27–30^. VOs provide a unique and ideal *in vitro* model to address these questions and resolve the controversies.

Three-dimensional retinal organoids (ROs) from PSCs have been established and evolved since 2011^31–35^, followed by organoids of the cornea^36–38^, ciliary body^39^, and lentoid body^40^. However, there have been no methods available for the generation of the vitreous body, the largest component of the human eyeball. Herein, using a stepwise differentiation, we generated the human vitreous organoids (VOs) from PSCs and identified their properties in terms of molecular signature, optical transparency, cell components, disease modeling and intraocular transplantability.

## Results

### Spatiotemporal analysis revealed fibroblasts as critical cellular orchestrators of vitreous morphogenesis

To delineate the spatiotemporal dynamics of vitreous morphogenesis, we conducted spatial transcriptomic profiling of human fetal ocular tissues across key developmental windows from post-conception weeks 8 to 15 (W8-W15). This approach revealed a dynamic redistribution of fibroblasts that orchestrates vitreous expansion (Fig. 1a). During early ocular development (W9), fibroblasts densely populated the vitreous core in both human (W9-W15) and murine (E11.5-E15.5) systems, exhibiting strong co-localization with Collagen I/II deposition (Fig. 1b-e) and expressing fibroblasts markers (FSP1, DCN; Fig. 1e-f). Strikingly, fibroblasts underwent strategic repositioning to a newly identified “growth edge” at the vitreous-retinal/lens interface (Fig. 1g), a region enriched in Collagen I/II/IV across species (human W15; mouse E16.5) (Fig. 1g and Extended Data Fig. 1a). Immunophenotyping confirmed fibroblast enrichment (MGP+) along this boundary, contrasting with hyalocyte markers (IBA1+/CD68+) in adjacent regions (Extended Data Fig. 1b-1c).

**Figure 1.**
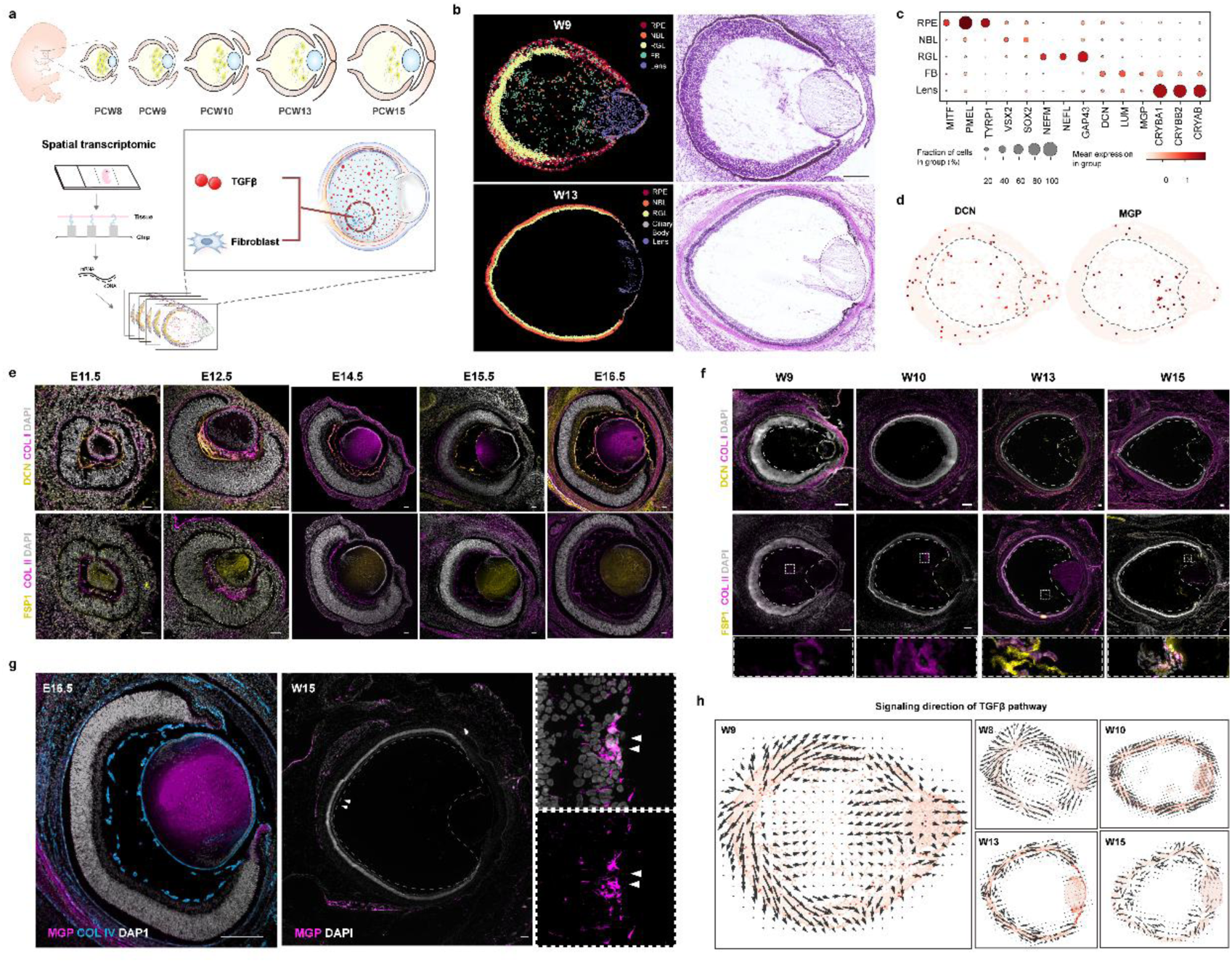
Spatiotemporal Coordination of Vitreous Fibroblast Dynamics and TGF-β Signaling During Embryonic Ocular Development. **(a)** Schematic diagram depicting the spatial transcriptomic workflow, sample preparation, and analytical outputs. **(b)** Spatial transcriptomics analysis of human embryonic eyes at weeks 9 (W9) and 13 (W13) post-conception reveals abundant fibroblasts within the vitreous at W9. Scale bars, 200 μm. **(c)** Cell cluster-specific marker gene expression profiles in the W9 human embryonic eye. **(d)** Spatial distribution of fibroblast-specific markers in the W9 human embryonic eye. **(e)** Immunostaining of murine embryonic eyes (E11.5-E16.5) demonstrates dense fibroblast populations (FSP1+, DCN+) and collagen fibers (Collagen I/II) within the developing vitreous. Scale bars, 50 μm. n = 2 biological replicates. **(f)** Temporal expression patterns of fibroblast markers (FSP1+, DCN+) and collagen fibers (Collagen I/II) in human embryonic eyes (W9-W15). Scale bars, 200 μm. **(g)** Late-stage fibroblast localization at the vitreous “growth edge” (vitreous-retinal and vitreous-lens interfaces) revealed by Collagen fibers and MGP immunostaining. Scale bars, 200 μm. (h) TGF-β signaling direction analysis during W8-W15: Early-stage (W8-W10) fibroblasts exhibit inward TGF-β pathway activation within the vitreous core.

Notably, pathway analysis pinpointed TGF-β signaling as the dominant regulatory axis, displaying stage-specific activation patterns that mirrored fibroblast redistribution (Fig. 1h). Early TGF-β hyperactivity (W9) coincided with fibroblast proliferation in the vitreous core, while its spatial restriction to the growth edge at later stages (W13-W15) aligned with matrix remodeling at the expansion frontier. In contrast, other pathways (WNT, VSFA, SPP1) showed minimal activity during vitreogenesis (Fig. 1h and Extended Data Fig. 1d), underscoring TGF-β’s unique role.

### Generation of vitreous organoids (VOs) from human pluripotent stem cells

TGFβ family members (TGFβ, TGFRs, BMPs and BMPRs) are highly expressed at the interactive surfaces of ocular tissues such as cornea, lens, vitreous, retina and RPE, and play important roles for mutual induction during development and retinal diseases^41–45^. TGFβ itself is also important for fibroblast proliferation, activation, transformation and matrix production^46,47^. We hypothesize that retina-vitreous crosstalk and fibroblasts are two critical drivers of vitreous development, and the addition of TGFβ to the retinal organoid induction medium^32,48^ can generate VOs.

Indeed, we successfully generated human VOs from all the 4 tested PSC lines, the BiPS, UiPS, H9, and CRX-tdTomato cell reporter lines, demonstrating the reproducibility and robustness of the protocol (Fig. 2a and Extended Data Fig. 2a-b). Interestingly, two types of human VOs could be distinguished based on their size and morphology (Fig. 2b-e). Type 1 VOs appeared opaque and smaller, whereas type 2 VOs were transparent and larger (Fig. 2c-e). By 7 months, the type 2 VOs could grow to more than 1 cm in diameter, whereas type 1 VOs could reach approximately 1 mm in diameter (Fig. 2b-f). The ratio of type 1 to type 2 organoids is approximately 79:21 in all cell lines (Fig. 2g).

**Figure 2.**
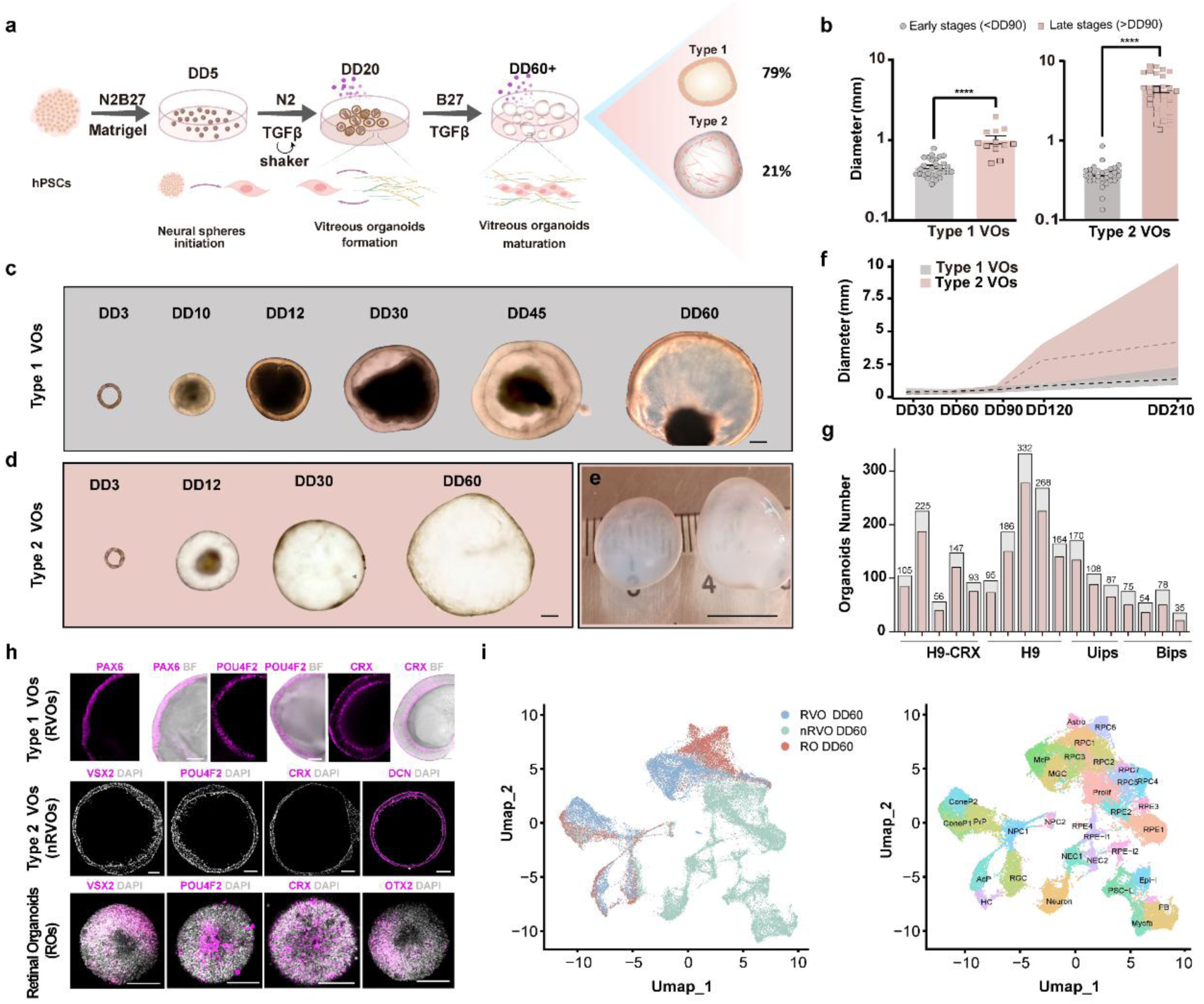
Generation of human vitreous organoids (VOs) from PSCs. **(a)** A schematic shows the procedure for VO differentiation at different stages. **(b)** The statistical calculations of the diameters of the type 1 and 2 VOs at early and late stages. ****, p < 0.0001. **(c-d)** Representative brightfield images for two types of VOs. Scale bars, 50μm. **(e)** Representative brightfield image of type 2 VOs at DD210. Scale bar, 1cm. **(f)** The presentation of diameters for two types of VOs at different timepoints. Dotted lines indicate the median of diameters. **(g)** Differentiation efficiency for four PSC lines, pink column shows the number of type 1 VOs. Grey column shows the total organoids in a batch. **(h)** Immunostaining Identification of two types of VOs and ROs. Scale bars, 200μm. n = 5 biological replicates. **(i)** Single cell sequencing analysis for RVOs, nRVOs and ROs at differentiation day 60.

To identify the two types of VOs, we used the whole-mount immunostaining to analyze the cell types at around differentiation day 60 (DD60). Since the mutual induction between retina and vitreous is beneficial for reciprocal development, and the “shell” of the type 1 VOs was stratified and appeared to be retina, we immunostained it for the retinal cell markers PAX6 (retinal progenitor cells, RPCs), CRX (photoreceptor precursors), POU4F2 (retinal ganglion cells, RGCs), SOX2 (RPCs) and OTX2 (RPCs), and all these markers are positive (Fig. 2h top panel and Extended Data Fig. 2c). Therefore, we referred to this type of VOs as the retinovitreous organoids (RVOs). Unlike RVOs and ROs, the type 2 VOs exhibited distinct molecular characteristics, absence of retinal lineage markers (VSX2, POU4F2, and CRX) alongside expression of fibroblast markers (DCN, VIM, KRT19) and co-expressed with Müller cell markers glutamine synthetase (GLUL) and S100B (Fig. 2h middle panel and Extended Data Fig. 2d), the shell of type 2 VOs did not stratify as retina. This type of VOs was termed as non-retinal vitreous organoids (nRVOs).

We performed single cell sequencing of DD60 ROs, RVOs, and nRVOs to comprehensively analyze their cellular components. Consistently, the RVOs and ROs were composed mainly of major retinal cell types (Fig 2i and Extended Data Fig. 2e). In contrast, the nRVOs had many non-retinal cell types, including fibroblasts, myofibroblasts, PSC-like cells, epithelial-like cells, and non-retinal neurons (Fig. 2i). The next issue is that the vitreous-specific proteins need to be further identified before RVOs and nRVOs can be claimed as vitreous organoids.

### Identification of characteristic vitreous gel proteins in VOs

The extracellular matrix (ECM) is responsible for stabilizing the vitreous gel structure^6,7^ and consists mainly of collagens (e.g., type II and type IV collagen) and non-collagenous structural proteins (e.g., fibrillin, GAPDH, tubulin), as well as some other macromolecules (e.g., fibronectin, laminin, agrin proteoglycans)^4,8^. To investigate the gel protein components of the VOs, we conducted the label-free proteomics for RVOs and nRVOs, or VOs that were depleted of the vitreous humor (VH, the liquid form of vitreous gel), and their VH (Fig. 3a).

**Figure 3.**
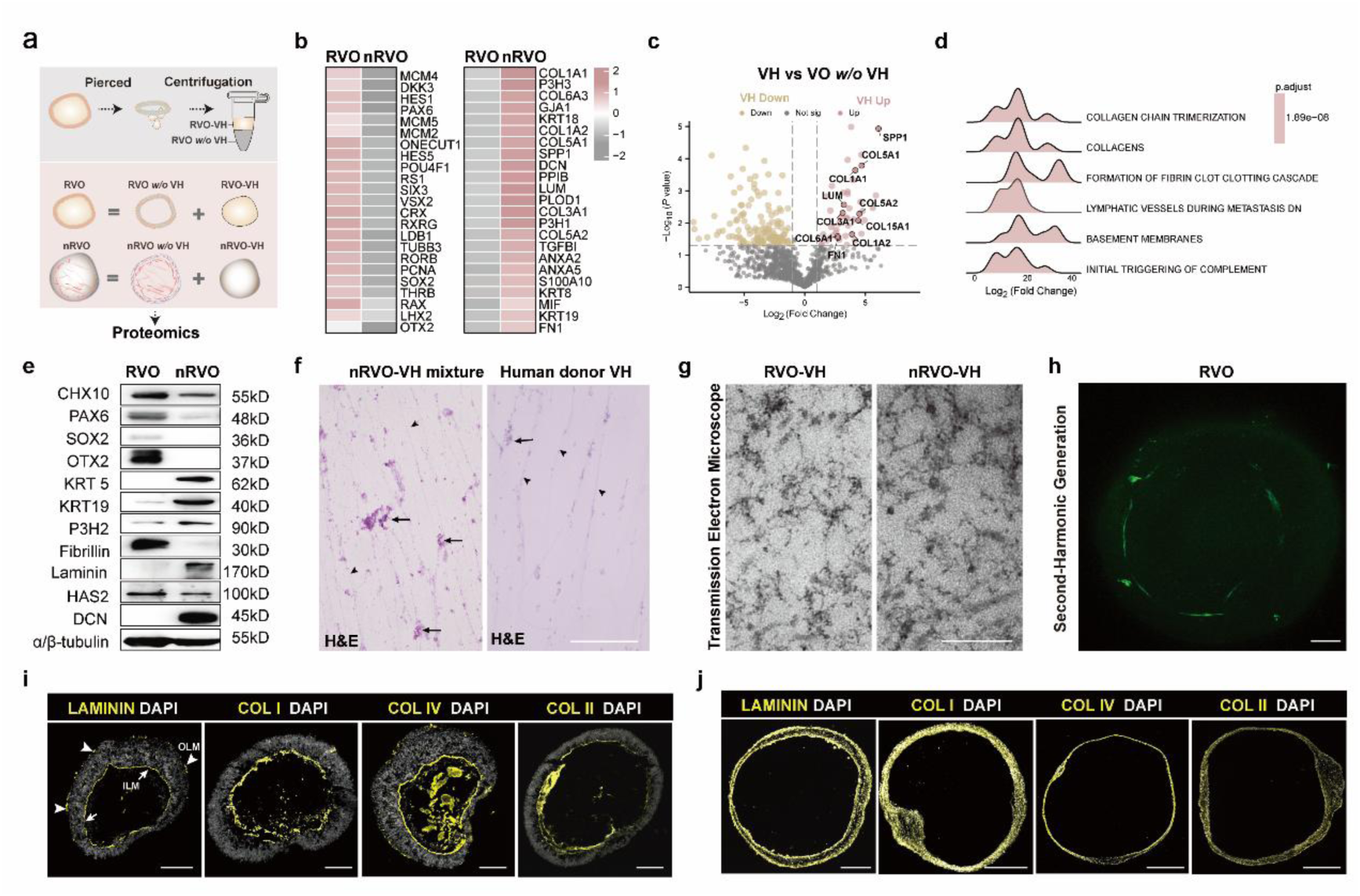
Characterization of structural protein components in RVOs and nRVOs. **(a)** A diagram illustrates the steps involved in sample preparation and sample compositions. **(b)** The heatmap based on label-free proteomics for RVOs and nRVOs showed the enriched proteins in each type of VOs. **(c)** The Volcano plot showed the differentially expressed proteins between the VH and VOs (depleted of VH). **(d)** Gene Set Enrichment Analysis (GSEA) result showed the enriched gene sets in the upregulated differentially expressed proteins (in e) in the VH group using the hallmark reference gene sets. **(e)** Western blotting results indicated that certain enriched proteins were expressed in RVO (n=20) and nRVO (n=4). **(f)** Transmission electron microscopy (TEM) image showed the fibrils in the VH of RVO and nRVO. Scale bar, 200nm. **(g)** H&E staining of the VH smears (from VH mixture of RVOs and nRVOs, or human donor VH) on slides showed the enriched collagen fibrils (indicated by arrowheads) and glycosaminoglycans (GAGs)/ proteoglycans (PGs)/ glycoproteins (GPs) (indicated by arrows). Scale bars, 200μm. **(h)** Collagens (green) were identified by the SHG imaging using a two-photon microscope. Scale bar, 100μm. n=3 biological replicates. **(i,j)** Immunostaining of RVOs (I) and nRVO (J) showed the expression of representative structural proteins LAMININ, COL I, IV and II arrows indicate inner limiting membrane (ILM)-like structures and arrowheads indicate outer limiting membrane (OLM)-like structures. Scale bar, 100μm in I and 500μm in J. n=3 biological replicates.

When comparing RVOs to nRVOs, proteomic analysis revealed that most retinal cell markers were lower in nRVOs, including RPCs markers (MCM4/5, DKK3, HES1/5, PAX6, PCNA, SIX3, VSX2, RAX, LHX2 and others), photoreceptor markers (CRX, RXRG, RORB, THRB), RGC markers (POU4F1, TUBB3), horizontal and amacrine precursor markers (ONECUT1), photoreceptor and bipolar markers (OTX2, RS1), except for Müller glial cell markers (GLUL, CLU, RLBP1), which were increased in nRVOs (Fig. 3b). Many of the vitreous collagens (COL1A1/2, COL3A1, COL5A1/2, COL6A3) and non-collagenous structural proteins (FN1, KRT8/18/19) were upregulated in the nRVOs (Fig. 3b). Notably, fibroblast-derived proteins (SPP1, DCN, PPIB, LUM, PLOD1, P3H1 and others) were enriched in nRVOs. When comparing the VH of nRVOs with VH-depleted nRVOs, differential expression analysis showed that the upregulated proteins in the VH of nRVOs were related to the formation of structural fibrils (Fig. 3c-d and Extended Data Fig. 3a-b), indicating that the vitreous structural proteins were enriched in the vitreous humor. Western blotting also conducted and identified these collages and non-collagenous structural proteins in RVOs, nRVOs, VH-depleted VOs, VH and human VH (Fig. 3e and Extended Data Fig. 3c-d).

Collagen fibrils are the characteristic structural features of the vitreous. The Hematoxylin-eosin (H&E) staining showed that abundant collagen fibers/fibrils, and glycosaminoglycans (GAGs), proteoglycans (PGs), and glycoproteins (GPs) supporting the scaffold of collagen fibrils^4^, were found in the VH of both VOs and the healthy human specimen (Fig. 3f). At a much higher resolution, the TEM image showed that the structural fibrils in the VOs (Fig. 3g) were comparable to those of the vitreous *in vivo*^8^. Second harmonic generation (SHG) is a specific label-free examination indicator of collagen fibrils. SHG imaging identified the coarse collagen fibers separating the VH from the retina shell, but was not sufficient to reveal the fine fibrils in the RVOs using two-photon microscopy (Fig. 3h and Extended Data Movie.1).

To identify the structural protein components of the vitreous and limiting membranes, we performed the immunostaining analysis of RVOs and nRVOs. In RVOs, laminin, the core basement membrane component, delineated the outer limiting membrane (OLM)-like and inner limiting membrane (ILM)-like structures (Fig. 3i), reproducing the OLM and ILM structures *in vivo*. The ILM is the vitreoretinal interface that separates the retina from the vitreous body^7^. The ILM in RVOs was also contributed by the structural proteins COL I, II and IV as *in vivo* (Fig. 3i and Extended Data Movie.2). Notably, COL I, II and IV were mostly enriched in the vitreous cortex, but were also present in the vitreous core (Fig. 3i), which was easily washed away during immunostaining. In nRVOs, laminin was expressed on the outer and inner shell surface (Fig. 3j). COL I, II and IV also contributed significantly to nRVOs (Fig. 3j). The results indicated that the structural proteins of the vitreous were expressed in the two types of VOs, whereas the RVOs were closer to the *in vivo* structural features of the retina-vitreous.

### Characterization of retinal tissue in RVOs

To compare the retinal tissues in RVOs and ROs, we focused on mature RVOs to investigate their retinal development. Because DD90 is a stage boundary of retinal organoid (RO) differentiation^49^, we consider the early stage to be the differentiation period before DD90 and the late stage to be after DD90.

Late-stage ROs have the laminated assembly and typical photoreceptors^31,32^. To verify whether the late-stage RVOs retained these retinal features, we cultivated the RVOs until DD240, when the photoreceptors were considered mature. Transmission electron microscopy (TEM) analysis of RVOs confirmed a laminated architecture (Fig. 4a) and the characteristic photoreceptor inner and outer segments (IS and OS) as in ROs (Fig. 4a-b). The photoreceptor OS is very fragile and easily detached during sample preparation. Nevertheless, some intact photoreceptors with the IS, connecting cilia (CC) and OS were identified (Fig. 4b). The ILM separating the retina from the vitreous was clearly visible (Fig. 4c). Interestingly, laminocytes, the vitreous-specific cells that predominantly align the posterior hyaloid membrane-ILM surface^50^ and maintain the adhesion *in vivo*^51,52^, were identified with characteristic cashew-shaped nuclei (Fig. 4d). Alternatively, scan electron microscopy showed that photoreceptor ISs were abundant (Fig. 4e) and some photoreceptors had intact IS, CC, and OS (Fig. 4f). These results confirmed the formation of a typical vitreoretinal surface and morphologically intact photoreceptors.

**Figure 4.**
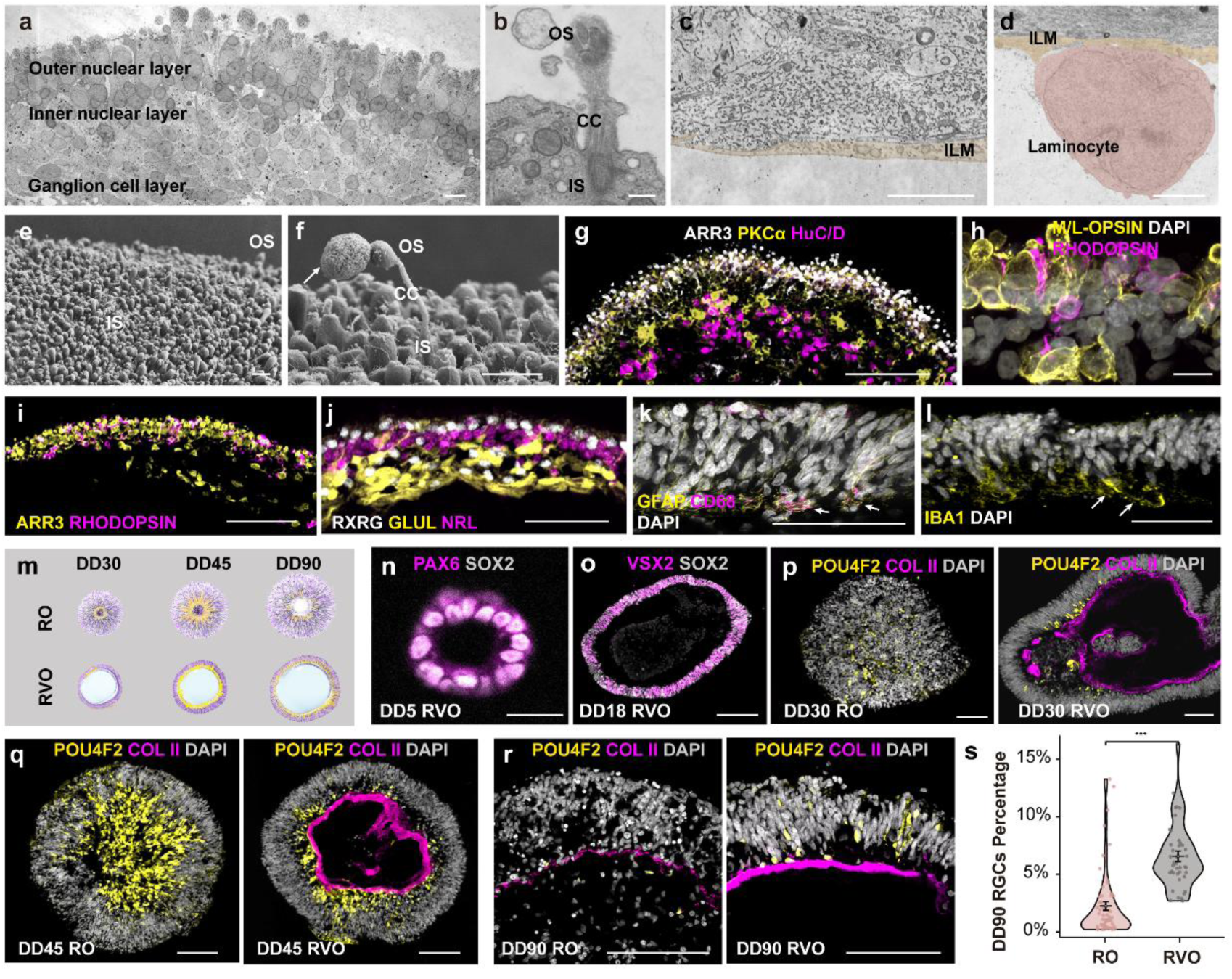
Structural and Cellular Characterization of RVOs. **(a)** The TEM images of DD240 RVOs showed the three-layered organization ILM (indicated in **c,d**), and photoreceptors with the intact inner segment (IS), connecting cilia (CC), and OS (in **b**). A typical laminocyte is shown with pseudopurple color in **d**. Scale bars, 5 μm. (E-F) Scanning electron microscopy (SEM) images showed the widespread IS on the surface of the DD240 RVOs. The OS was seen sporadically because most of it was sheared off during the specimen preparation. Scale bars, 10μm. **(g-l)** Fluorescence immunostaining showed the formation of cones (ARR3+, M/L opsin+ in g-l), rods (RHODOPSIN+ in h-i, NRL+ in j), bipolar cells (PCKα+ in g), RGCs (HuC/D+ in g), and Müller cells (GLUL+ in j) in DD240 RVOs. **(k)** GFAP+; CD68+ double positive laminocytes at the vitreoretinal interface. Arrows point to double immunopositive cells. **(l)** IBA1+ hyalocytes at the vitreoretinal interface. Arrows indicate IBA1+ cells. Scale bars, 50 μm. n=4 biological replicates. **(m)** Schematic illustration of RGC development in ROs vs. RVOs (days 30-90) **(n)** Immunostaining for neural spheres of RVO at differentiation day 5. Scale bars, 20 μm. n=10 biological replicates. **(o)** Representative images of retinal progenitors in RVO at differentiation day 18. Scale bars, 100 μm. n=5 biological replicates. **(p-r)** Comparative immunofluorescence for collagen II and RGCs in ROs vs. RVOs (days 30-90). Scale bars, 100 μm. **(s)** Quantitative comparison of RGC numbers at day 90 between ROs (62 sections/5 organoids) and RVOs (40 sections/4 organoids).

We next examined the major retinal cell types in DD240 RVOs by immunostaining. Antibodies against the cone markers (ARR3, M/L-OPSIN, RXRG) and the rod marker (RHODOPSIN, NRL) revealed the formation of M/L cones (ARR3+; RXRG+; M/L-OPSIN+), S cones (ARR3+; RXRG+; M/L- OPSIN-), and rods (RHODOPSIN+; NRL+; ARR3-) (Fig. 4g-j). We also identified the interneuron rod bipolar cells (PKCα+) and the efferent neuron RGCs (HuC/D+; RXRG+) (Fig. 4g and j). RXRG labels both cones in the outer nuclear layer and RGCs in the RGC layer (Fig. 4j), distinguished by their laminar position. The Müller cell marker glutathione synthetase (GLUL) confirmed their presence (Fig. 4k). More importantly, the laminocytes (GFAP+; CD68+) and hyalocytes (IBA1+), two cell types important for vitreous physiology^53^, were found adhering to the vitreoretinal surface (Fig. 4k-l), providing strong evidence for the formation of a vitreoretinal surface resembling the *in vivo* structure.

A significant challenge of contemporary RO technology is the inability of RGCs to survive, rendering ROs a poor tool to study RGC development and diseases^54^. RGCs are efferent neurons that integrate visual signals from the retina and send them to the brain^55^. Loss and dysfunction of RGCs will result in glaucoma, the leading cause of irreversible blindness worldwide. Unfortunately, studies of RO-derived RGCs have been limited in the early stages, typically DD30-90, because RGCs are gradually lost and almost disappear around DD120^54,56,57^. In both ROs and RVOs, RGCs were born around DD30 and enriched at DD45 (Fig. 4m-q). A significant proportion of RGCs survived to DD90 in RVOs (Fig. 4r). In contrast, only a limited number of RGCs can be detected in the inner layer of DD90 ROs (Fig. 4r-s). The reasons for the enhanced survival of RGCs in the RVOs remain unclear; however, we speculate that the vitreous may provide a form of support.

To evaluate the physiology of photoreceptor cells in DD220 RVOs, we performed patch-clamping using whole-cell recording. Photoreceptors appeared brighter in the outer layer under the microscope (Extended Data Fig. 4a). To determine if ion channels were functional, we measured the hyperpolarization-activated current (I*_h_* current), the voltage-gated sodium current (Na^+^ current), and the membrane voltage responses. Most photoreceptors in 3 RVOs exhibited significant inward I*_h_* currents under hyperpolarized conditions (Extended Data Fig. 4b). And 18 out of 42 photoreceptors displayed Na^+^ currents under voltage-clamp configuration (Extended Data Fig. 4c). We also observed Na^+^ spike-like responses under current-clamp recordings and found that photoreceptors with large Na^+^ currents exhibited spike-like responses (Extended Data Fig. 4c). Additionally, we assessed fundamental membrane properties such as resting membrane potential, membrane capacitance, and membrane resistance (Extended Data Fig. 4d-e). All these results are consistent with previous studies in human or non-human primate mature photoreceptors^58,59^.

Taken together, these data suggest that the retinal tissue in RVOs is similar to that in ROs, but has the advantage of supporting longer RGC survival, and that RVOs are comparable to the retina-vitreous *in vivo* in several aspects, including the retina and vitreous and the ILM interface that separates them.

### Cellular origin of ECM components in the vitreous of organoids

The cellular origin of vitreous macromolecules during development is still under debate. Considerable early evidence from RNA in situ hybridization and immunostaining suggests that surface ectoderm and fibroblasts from the developing ciliary body are the dominant origin of vitreous proteins^7^. Others argue that the fibroblasts from the hyaloid vessels and lens vesicle, and Müller cells from the retina are the contributors^8^. We collectively examined all the cell types in DD60 nRVOs, RVOs, and ROs, and found that fibroblasts were abundant in nRVOs, some in RVOs, but not in ROs (Fig. 5a), suggesting that fibroblasts were not originated from lens or ciliary bodies *in vitro*.

**Figure 5.**
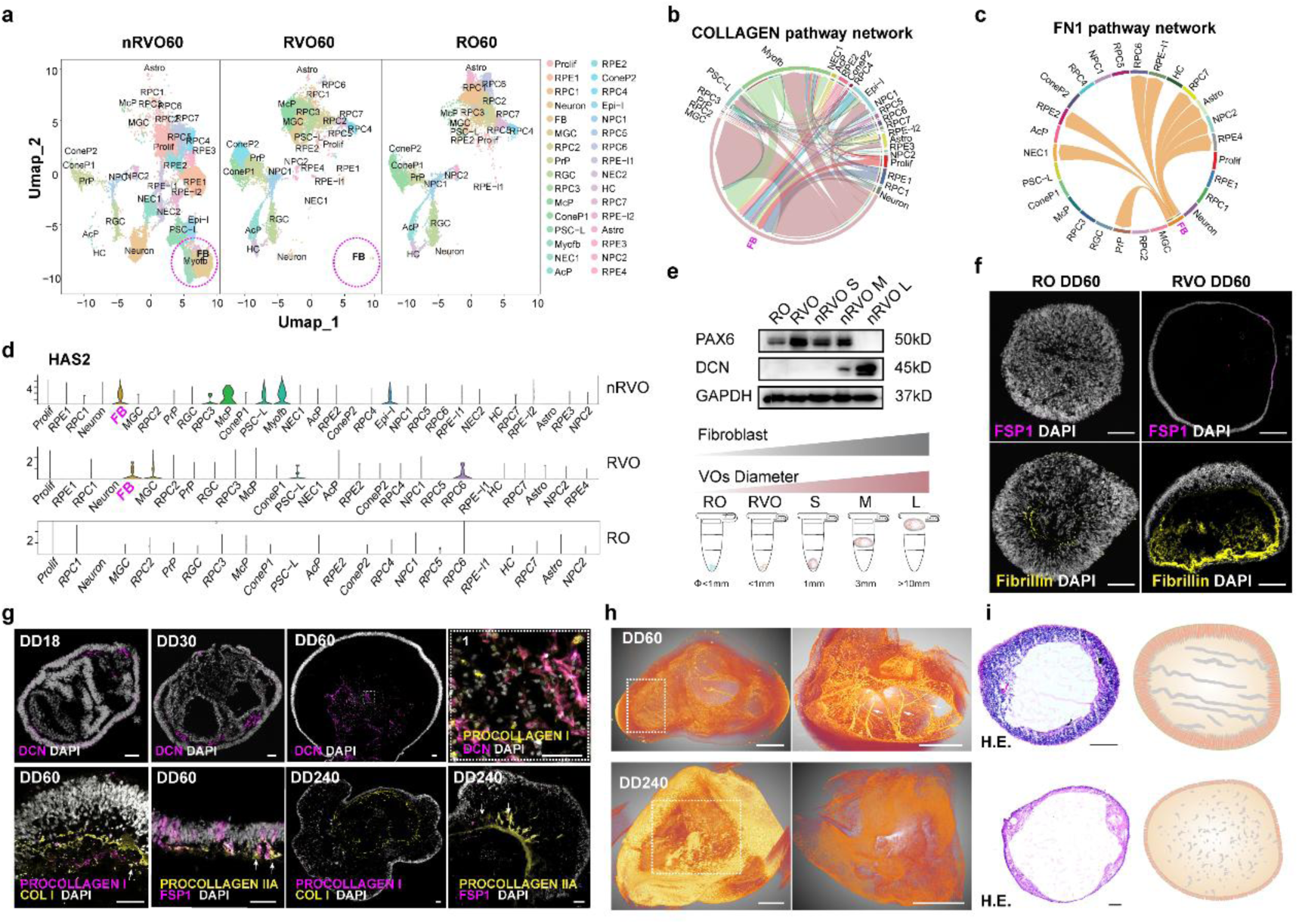
Dynamic changes in fibroblast function and matrix remodeling in vitreous organoids. **(a)** Single-cell analysis of DD60 nRVO, RVO, and RO. Purple ellipses highlight fibroblasts. **(b)** Collagen fiber signaling pathway in nRVO, primarily associated with fibroblast-mediated synthesis and secretion. **(c)** FN1 signaling pathway in RVO, exclusively produced by fibroblasts. **(d)** Expression patterns of HAS2 (hyaluronic acid synthase gene) across cell types. Negligible expression in RO, with fibroblast-specific expression in both nRVO and RVO. **(e)** Western blot analysis of fibroblast markers in nRVO, RVO, and RO. Fibroblast content correlates with organoid size. **(f)** Immunofluorescence staining demonstrates significantly higher fibroblast (FSP1+) and matrix (Fibrillin+) content in RVO compared to RO. Scale bars, 100μm. n=3 biological replicates. **(g)** DCN immunostaining reveals progressive fibroblast accumulation from D18 to D60 in RVO. Fibroblast (DCN+/FSP1+) co-localization with both procollagen I and II as well as the procollagen I with collage I. Scale bars, 50μm. n=5 biological replicates. **(h)** Micro-CT imaging shows intact fiber networks in DD60 RVO versus fragmented punctate fibers in DD240 RVO. Scale bars, 200μm. n=2 biological replicates. **(i)** H&E staining and schematic illustration confirm temporal fiber integrity changes: intact fibers in early stages versus fragmented patterns in later stages. Scale bars, 200μm. n=3 biological replicates.

As RVOs are closer to the retinovitreous *in vivo*, they were used to investigate vitreous development. At the transcriptional level, fibroblasts expressed unique (*COL1A1, COL1A2, COL3A1, COL5A2, COL6A3, COL12A1, COL14A1, COL16A1, COL21A1, PCOLCE, and PLOD3*) and predominant (*COL5A1, COL6A2, COL27A1, and PLOD1*) genes involved in collagen anabolism (Extended Data Fig. 5). CellChat analysis of cell-cell interactions showed that fibroblasts (and derived myofibroblasts) interact with almost all cell types and are the predominant signaling senders in the collagen pathway network (Fig. 5b). For non-collagen protein genes, CellChat showed that fibroblasts are the sole signaling senders of the FN1 pathway network (Fig. 5c). In human vitreous, hyaluronan, the major polysaccharide component, fills the spaces between the collagen fibrils^4^. The mRNA encoding the enzyme that synthesizes hyaluronan primarily in the retinovitreous, termed the hyaluronan synthase 2 (HAS2), was examined in cell types of the DD60 organoids. *HAS2* mRNA levels were found to be minimal in RO60 cells, yet they were detected in fibroblasts (and myofibroblasts) and several other cell types within RVOs and nRVOs (Fig. 5d), suggesting that fibroblasts produce hyaluronan and may also induce other cell types to produce hyaluronan via a paracrine manner. These results support that fibroblasts may play a pivotal role during vitreous formation.

To further validate their role, we performed immunostaining and Western blotting to examine fibroblast cell markers and secreted products. We hypothesize that the number of fibroblasts is positively associated with the diameter of VOs (Fig. 5e). Indeed, the protein level of the fibroblast marker DECORIN (DCN) was low in medium-sized nRVOs, very high in large nRVOs, but not detectable in ROs, RVOs, or small nRVOs (Fig. 5e). Surprisingly, the retinal cell marker PAX6 was drastically downregulated in large nRVOs. Consistent with the scRNA-seq results, FSP1 (S100A4), a fibroblast cell marker, was immunopositive in RVOs but not in ROs (Fig. 5f). The non-collagenous structural protein FIBRILLIN was strongly expressed in RVOs, but was very restricted in ROs (Fig. 5f). At early differentiation stages (DD18 and 30), DCN+ fibroblasts were found at the emerging retina-vitreous interface (Fig. 5g). Procollagen chains are an indicator of collagen synthesis^60,61^. In DD60 RVOs, a significant number of PROCOLLAGEN I and IIA particles were found to be colocalized with or in close proximity to the fibroblast markers DCN, FSP1, and COL I, suggesting that these particles were derived from fibroblasts (Fig. 5g). In mature DD240 RVOs, some FSP+ fibroblasts still produced PROCOLLAGEN IIA (Fig. 5g), suggesting that these cells continued to synthesize procollagens to maintain vitreous structure. Collectively, these results suggest that the TGFβ formula induced human PSCs to generate fibroblasts, and that fibroblasts were the major contributors to vitreous components including collagen and non-collagen proteins and polysaccharides.

### Dynamic ECM remodeling and cell apoptosis during RVO development

Dynamic ECM remodeling and gel liquefaction are hallmarks of vitreous development and aging *in vivo,* although the mechanisms are poorly understood^8^. To determine the spatiotemporal dynamics of the macromolecular scaffold in the vitreous, we examined the RVOs at various developmental stages. As indicated by MicroCT and H&E staining, the DD60 RVOs formed a coarse fibrillar ECM network, with fibrils perpendicularly inserted into the ILM, as previously observed *in vivo*^62^ (Fig. 5h-i). At DD240, however, the fibrils became thinner but denser, particularly in the vitreous cortex (Fig. 5h-i), suggesting extensive ECM remodeling. Matrix metalloproteinases (MMPs) have been shown to play an important role in ECM breakdown and vitreous remodeling^8^. The proteomic analysis revealed the expression of certain MMPs in the VOs and VH, suggesting a conserved ECM remodeling process during VO development *in vitro* (Extended Data Fig. 6g).

Programmed cell death (PCD) is imperative for tissue development; however, its role in vitreous development remains to be elucidated. The truth that a limited number of cells survive in the mature vitreous suggests the potential for apoptosis during this process. To investigate this, we performed a TUNEL assay, which revealed that a significant number of cells underwent apoptosis in DD60 RVOs, particularly in the vitreous cortex. The TUNEL- positive signals increased at DD90, but were rare at DD240 (Extended Data Fig. 6a-d). We also examined the active apoptosis effector, cleaved caspase 3 (CL.CASP3), in the retina of RVOs. Correlating with TUNEL staining, CL.CASP3 signals were found in some cells at DD60, peaked at DD90, but almost disappeared at DD240 (Extended Data Fig. 6e-h). The presence of characteristic apoptotic features, including cell shrinkage, chromatin condensation and apoptotic bodies, was further confirmed by TEM in the cells of the VOs (Extended Data Fig. 6f). In the proteomic data, the apoptosis-related effectors (CASP3, CASP6, CASP7) and regulators (BCL2, BAX) were enriched in the retina of RVOs (Extended Data Fig. 6g). These findings provide substantial evidence that cell apoptosis is an essential process in the vitreous formation at early developmental stages (Extended Data Fig. 6h).

### Proof-of-concept transplantation of RVOs and vitreous humor in animal model

One of the important potential applications of VOs is vitreous regeneration following vitrectomy. To assess the transplantability of VOs, a vitrectomy was performed to remove a portion of the vitreous (Extended Data Fig. 7a-b), and then equal volumes of nRVOs or RVOs were transplanted into the rabbits. After two weeks, the rabbits were euthanized, and the eyeballs were examined using H&E and DAPI staining. A comparison of the eyes that received nRVOs (Extended Data Fig. 7d) with the non-transplanted eyes (Extended Data Fig. 7c) revealed the presence of severe proliferative vitreoretinopathy (PVR). This is likely attributable to Müller and stem cell proliferation in the nRVOs, which have been identified as a trigger for PVR^63^. Conversely, the eye transplanted with RVOs exhibited no indications of PVR (Extended Data Fig. 7e), thereby suggesting that RVOs may function as vitreous substitutes. However, it should be noted that the transplantation of whole RVOs necessitates a substantial retinal transection, a procedure that may potentially result in retinal detachment (Extended Data Fig. 7d-e). Consequently, the utilization of VH as an alternative to RVOs could be a more pragmatic and effective approach for transplantation.

To create a vitreous liquefaction disease model, 5 units of hyaluronidase were injected into the rabbit vitreous (Extended Data Fig. 7f). After one week, the liquefied vitreous was replaced with control PBS, or VH from RVOs, or healthy human VH from an elderly donor. Two weeks later, the rabbit vitreous was examined by B-scan ultrasonography (Extended Data Fig. 7f). The results showed that all grafts did not alter the vitreous transparency, indicating the absence of PVR (Extended Data Fig. 7g). Notably, B-scan ultrasound images revealed the presence of floaters in the vitreous of all graft groups (PBS, organoid VH, and donor human VH) (Extended Data Fig. 7g), which is a common symptom of vitreous liquefaction. However, the organoid VH transplantation exhibited a more pronounced alleviation of these floaters compared to PBS or donor human VH transplantation (Extended Data Fig. 7g). This observation suggests that the organoid VH, which is readily available, is compatible with *in vivo* VH and demonstrates its potential for clinical vitreous replacement.

### RVOs recapitulated familial vitreous amyloidosis disease caused by a common *TTR* mutation

We wonder if RVOs could be used as vitreous disease models to recapitulate symptoms or phenotypes observed in patients. Familial vitreous amyloidosis (FVA) is characterized by the accumulation of the amyloid transthyretin (ATTR) fibrils in the vitreous, typically caused by the autosomal dominant mutations in the *TTR* gene^13,14^. However, the origin of ATTR remains a subject of debate. One hypothesis suggests that the liver is the primary site of TTR synthesis, with TTR being transported to the vitreous via blood circulation to form amyloid deposits. Alternatively, ATTR may originate from adjacent tissues such as the RPE and retina (Fig. 6a). We examined the cellular sources of TTR by assessing the expression of *TTR* mRNAs. Consistent with prior reports in rat^64^, TTR mRNA was predominantly detected in developing human RPE cells (Extended Data Fig. 8a); in addition, it was also expressed at low levels in many other retinal cell types of DD60 ROs, RVOs and nRVOs (Extended Data Fig. 8a). Additionally, we found that *TTR* is among the top most highly expressed variable genes in nRVOs (Extended Data Fig. 8b). Furthermore, proteomics data showed that TTR protein was highly enriched in the VH of RVOs and nRVOs, rather than in the retinas (Fig. 6b). These data suggest that the biosynthesis of TTR is conserved in vertebrate retinas and that retinal cells, in addition to hepatic cells, are capable of synthesizing TTR.

**Figure 6.**
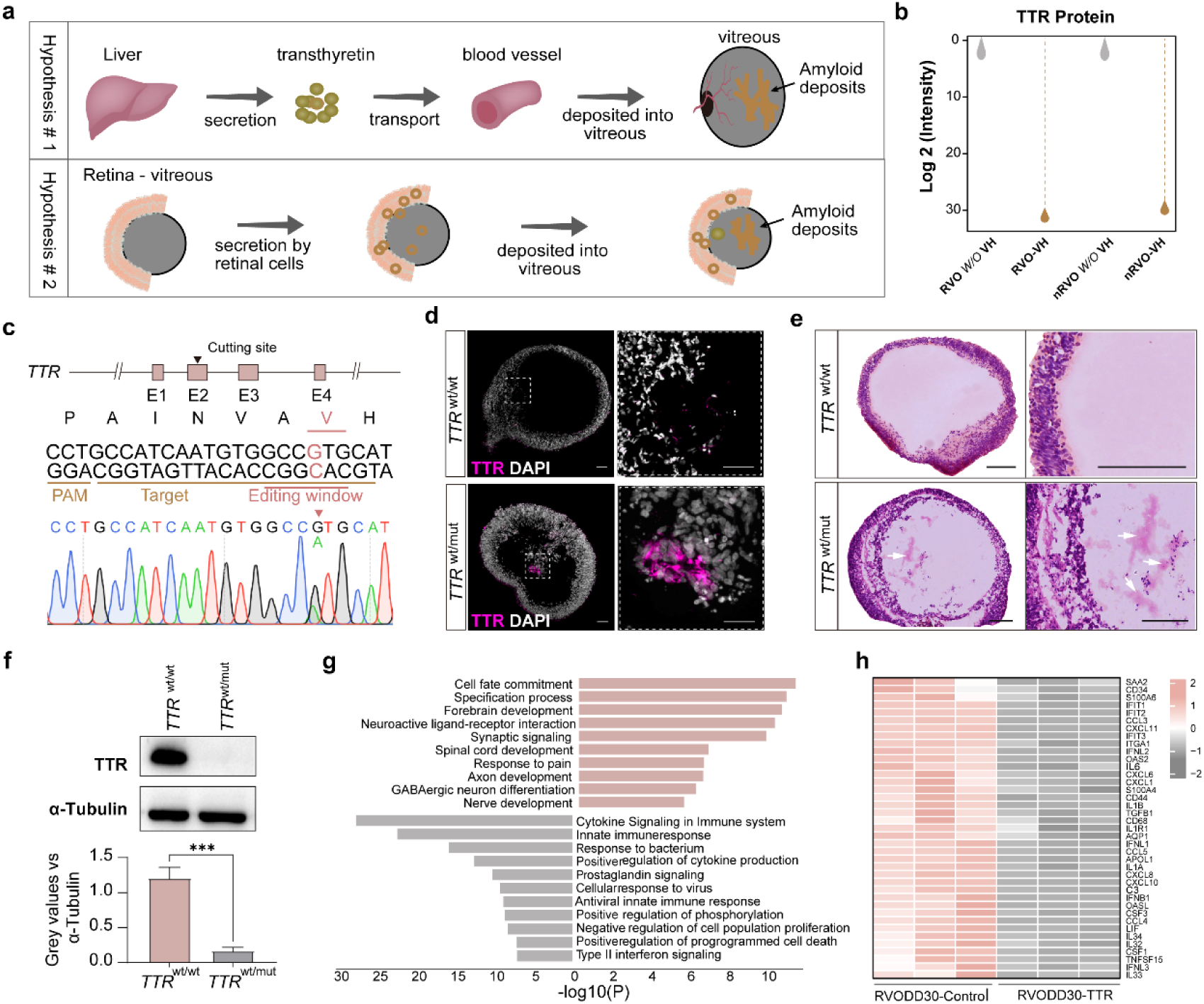
Mechanisms of TTR mutation-induced familial vitreous amyloidosis disease in vitreous organoid models. **(a)** illustration of two hypotheses that the vitreous amyloid deposits in familial vitreous amyloidosis disease originate from the liver or adjacent tissues. **(b)** Proteomic sequencing confirmed TTR protein was exclusively detected in vitreous humor, absent in humor-depleted vitreous organoids. **(c)** Illustration of the TTR genomic structure, design of the TTR V30M base editing strategy, and verification of the edited sequence by Sanger sequencing. **(d)** TTR protein exhibits sparse localization within wild-type RVOs, while TTR-mutant RVOs show prominent TTR aggregates (arrowheads) in vitreous-like compartments. Scale bars, 50μm. n = 3 biological replicates. **(e)** Congo Red staining demonstrated amyloid deposits (Congo Red+) in TTR-mutant RVO. The arrow indicates the Cango red-positive gel-like fibril deposits in the vitreous. Scale bars, 200μm. n=5 biological replicates. **(f)** Western blot analysis of TTR monomer levels in wild-type and TTR-mutant samples, normalized to α-tubulin (loading control). Graph quantifying TTR/α-tubulin ratios, demonstrating significant reduced monomeric TTR levels in mutants compared to wild-type. ***, p < 0.001. **(g-h)** RNA-seq analysis showed significant downregulation of immune-inflammatory pathways (g) and corresponding gene expression in TTR mutants versus controls (h).

The dominant TTR V30M mutation is the primary cause of FVA^65^, and to model FVA, a heterozygous *TTR* (V30M) mutation was introduced into the H9- CRX-tdTomato line using the base editing technology (Fig. 6c). The accumulation of the TTR amyloid can be detected by the Congo red staining and immunostaining of TTR proteins (Extended Data Fig. 8c). After differentiation of *TTR*-mutated PSCs into RVOs, the Congo red-positive gel-like amyloid deposits and TTR proteins could be identified by DD60 RVOs (Fig. 6d-e), Western blotting results show a decline of TTR monomer levels in TTR-mutant RVOs at DD60, which may be caused by TTR aggregation into stable polymeric forms (Fig. 6f). Interestingly, TGFB1 also displayed a decrease in TTR mutant RVOs (Extended Data Fig. 8d). These results suggesting that retina itself is sufficient for vitreous ATTR deposits in patients.

To further explore the molecular changes underlying the *TTR* mutation in RVOs, we performed bulk RNA sequencing. This analysis revealed that the differentially expressed genes (DEGs) in TTR-mutated RVOs compared to wildtype ones were primarily enriched in neural development pathways, including cell fate commitment, specification process, forebrain development, synaptic signaling, and related processes (Fig. 6g). Conversely, the downregulated DEGs were predominantly enriched in the immune-related pathways, including the cytokine signaling in immune system, innate immune response, and prostaglandin signaling (Fig. 6g-h), which is consistent with previous reports that wildtype but not mutated TTR could stimulate proinflammatory responses in Schwann cells and macrophages^66,67^. Collectively, these results underscore the utility of RVOs as a valuable model for studying vitreous and vitreoretinal diseases.

## Discussion

### *In vivo* and *in vitro* vitreous development

The human vitreous constitutes approximately two-thirds of the total volume of the eyeball, making it the most substantial component of the eye. However, its significance has often been neglected, in part due to the difficulty of obtaining human tissue. A comparison of human and mouse vitreous development reveals several common and conserved features. For instance, both species exhibit similar processes such as the formation and subsequent degeneration of the hyaloid vasculature, vitreous remodeling, and other processes. Despite the substantial structural disparities between these two components, the mouse vitreous can offer significant insights into the complex developmental processes of the human vitreous.

Embryonic vitreous development includes three stages, the primary, secondary, and tertiary vitreous^8^. The primary vitreous is formed along with the hyaloid vasculature^68^. With gradual remodeling, the primary vitreous transitions to the secondary vitreous, which is the definitive (adult) vitreous^69^. The tertiary vitreous is associated with the lens zonules that form at the developing ciliary body^8,68^. Our protocol for vitreous development *in vitro* does not include such stages.

This is the first time to use spatial transcriptomics to study human vitreous development. In human and mouse vitreous development, we discovered a vitreous ‘growth edge’ close to the retina-vitreous and lens- vitreous interfaces across all developmental stages, where fibroblasts persistently produce vitreous structural proteins and hyaluronan. To some extent, the observation of the growth edge could perfectly explain how the developing vitreous keep expanding. Further analysis of cellular interaction network revealed that the TGFβ signaling is an important pathway regulating vitreous development. Take advantage of the discovery, we have robustly generated human VOs from ESCs and iPSCs. Amazingly, the volume of VOs is comparable to that of human neonates, and they are transparent in nature. Two types of VOs, RVOs and nRVOs, were identified based on their morphology and cell types. Meanwhile, the vitreous humor (VH) of VOs is comparable to the *in vivo* VH not only in the transparency but also in biological characteristics and molecular composition.

*In vitro* RVOs provide a simplified system for studying vitreous development. RVOs have a similar retina-vitreous assembly with an ILM-like interface, but do not have the hyaloid vasculature, lens, iris, or ciliary body. This abridged system offers the advantage of eliminating the confounding effects of the vasculature, lens, and ciliary body on vitreous development. As evidenced from animals with avascular vitreous, such as avian, the hyaloid vasculature appears to be dispensable for vitreous development. However, in humans or animals with hyaloid vasculature, the vasculature might be vital for transporting the necessary nutrients and oxygen for expanding the vitreous to a greater volume. Additionally, the vasculature serves as a conduit for the entry of fibroblasts and myeloid cells into the developing vitreous.

To the best of our knowledge, the nRVOs represent the largest organoid in dish to date. Without the vasculature, the nutrient supplement is entirely dependent on diffusion of the culture medium, a process that imposes constraints on nRVOs’ growth and may introduce a stressful microenvironment within the organoids. As an indicator of stress in nRVOs, a notable proportion of fibroblasts transited into myofibroblasts. Future studies could explore measures to alleviate these constraints and enhance the system.

### The cellular contribution of vitreous macromolecules in vitreous development

The fibrillar meshwork in the vitreous is essentially composed of the ECM including proteins (mainly collagens) and other macromolecules. However, the cellular contribution of the macromolecules is still controversial. Studies in chick and mouse models, as well as human specimens, suggest that the ciliary body, lens, hyaloid vessels, and presumptive retina are the major sources of the vitreous collagens^4,7,8,29,70–73^.

Our results showed that fibroblasts directly contribute to many collagen and non-collagen proteins as well as the polysaccharide hyaluronan. Some of them are fibroblast-exclusive, including *COL1A1, COL1A2, COL3A1, COL5A2, COL6A3, COL12A1, COL14A1, COL16A1, COL21A1, PCOLCE, and PLOD3, a*mong which COL1A was considered a specific marker for fibroblasts in many tissues. Fibroblasts also instruct other cell types to produce fibrillar components. For instance, COL II is the most abundant collagen in human vitreous; although scRNA-seq data showed that fibroblasts were probably not the major contributors of COL II, and only those retinal cells adjacent to fibroblasts were capable of producing PROCOLLAGEN II in RVOs, a phenomenon attributed to positional effect. Fibroblasts are also the unique source of the FN1 signaling pathway, a mediator of positional effect. Zhang *et al.* reported that FN1 could stimulate chondrocytes to produce COL II and other ECMs via the TGFβ/PI3K/AKT pathway^74^. Whether FN1 has a similar role in retinovitreous warrants further investigation. We also found that the size of VOs was correlated with the number of fibroblast cells, demonstrating the importance of fibroblasts in vitreous development.

Taken together, our results strongly suggest that fibroblasts are the pivotal cells in vitreous development.

### Cell death and vitreous remodeling in human vitreous development

As one of the controlled programmed cell death pathways, apoptosis is essential to eliminate unwanted cells to ensure proper tissue structure during development^75,76^. The embryonic primary vitreous contains numerous cells but gradually decreases, apparently accompanied by cell loss due to cell death. During remodeling, the hyaloid vasculature regresses and vascular cells undergo cell apoptosis^77^. Here, we found in our *in vitro* model that a large amount of cell apoptosis could be identified in the early stage vitreous. Since there is no hyaloid vasculature in VOs, it is likely that apoptosis is not directly induced by regression of the vasculature as *in vivo,* but rather by hypoxic stress and/or low nutrient supply and/or other environmental factors. Consistent with the notion of environmental stress, fibroblasts (and myofibroblasts) expressed up to tenfold the TLR4 gene comparing to those in ROs (data not shown), which likely activates the DAMP signaling in fibroblasts and promotes ECM degradation by secreting remodeling enzymes such as MMPs^78^.

### Vitreous organoids for vitreous replacement after vitrectomy

Vitrectomy is a prevalent treatment modality employed in the management of vitreoretinal diseases. Given the non-regenerative nature of the human vitreous, artificial vitreous substitutes are inevitably required after vitrectomy. However, the existing vitreous substitutes, including inert gas, silicone oil, and hydrogels, have significant deficiencies in structural and, more importantly, functional alignment with the natural vitreous. These substitutes are incapable of sustaining the low oxygen tension and homeostasis within the vitreous, resulting in various postoperative complications such as cataract and glaucoma^25,79^. Consequently, VOs are emerging as an ideal post-vitrectomy substitutes with its comparable biophysiological and biochemical properties to endogenous vitreous. Our proof-of-concept transplantation experiments have shown that, when grafted into a rabbit model of vitreous liquefaction, RVO-derived VH exhibited robust safety comparable to healthy human vitreous and could effectively repair the vitreous damage. Despite the need for further refinement of the surgical procedure, the unlimited availability and the demonstrated transplantability of organoid VH make it a promising vitreous replacement to restore vitreous function.

### Vitreous organoids as *in vitro* models for basic and translational research

Organoids of retina, brain, liver, and others have been widely used in the basic and translational studies^80–82^. During VO development, we noticed that the fibrillar morphology and distribution vary with time, a phenotype called vitreous remodeling, which is considered as a physiological change with development and aging rather than a disease. However, how vitreous remodeling is initiated and progressed is unknown, which is important for understanding and treating various vitreoretinal diseases. The availability of unlimited VOs provides a new platform to uncover the mechanism of vitreous remodeling.

Familial vitreous amyloidosis (FVA) is an autosomal dominant genetic disease that causes progressive vision loss. We constructed the FVA disease model by introducing the *TTR* mutations into the RVOs using single-base editing technology, and successfully reproduced the typical gel-like amyloid accumulation in the vitreous. Our results prove that retina itself is sufficient for vitreous amyloid deposit; however, we can’t rule out the possibility that the liver also contributes to the vitreous deposit *in vivo*, which may accelerate the disease progression. We further confirmed that downregulation of immune-related pathways in the TTR-mutated RVOs, in agreement with previous reports.

The size of the vitreous body may determine the diameter of the eye during development and its regulatory mechanism is unknown. VOs may serve as a valuable tool to elucidate the regulation of human eye size and the pathogenesis of related diseases such as myopia and microphthalmia^9,11,12^.

In summary, we have established an effective and robust method to generate VOs from human iPSC and ESC cells. This advancement holds great promise for future studies of vitreous development and diseases, as well as for future regenerative therapies to treat a wide range of eye disorders, including vitreous floaters, macular holes, and other vitreoretinal interface-related diseases.

## Author contributions

Xiao Zhang performed the experiments, collected and interpreted the data, prepared the figures and wrote the manuscript. Kangxin Jin supervised the project, analyzed and interpreted the data, performed bioinformatic analysis and wrote the manuscript. Xiao Lin performed the experiments and revised the manuscript. Hai-Long He carried out transplantation. Mingxia Du performed the differentiation experiments. Fanying Zeng, Ping Liu collected and prepared the human embryo samples. Wen Wang performed the bioinformatic analysis. Yue Zhang performed the electrophysiologic analysis. Dongdong Zhao and Changhao Bi designed and performed the single base editing of the cell lines. Zi-Bing Jin conceived the concept, designed and supervised the project, critically revised the manuscript, and provided financial support. All authors approved the submission.

## Data availability

The RNA-seq data will be uploaded to Genome Sequence Archive with the access number HRA013489. Other data will be available upon reasonable request to the corresponding author.

## Declare of interest

A patent based on this project is pending. All authors declare that they have no competing interest.

## Acknowledgement

We thank Mr. Zhibao Zhang for technical support of Congo-red staining, Zhuo-Lin Liu, Shu-Ning Sun, Lingzi Wu and Jing Yang for performing the biochemical and immunostaining experiments, Yi-Xin Liu and Xiaowei Wang for revising the manuscript, Yan-Ping Yu for collecting human vitreous. This project was supported by fundings from the National Natural Science Foundation of China (82125007,92368206), Beijing Natural Science Foundation (Z200014, 5242005), Beijing Municipal Public Welfare Development and Reform Pilot Project for Medical Research Institutes (PWD&RPP-MRI, JYY2023-6) and the Beijing Hospitals Authority Clinical Medicine Development of Special Funding Support (DFL20220202, ZLRK202503), Excellent Clinical Research Program (BRWEP2024W172050100).

