## Supplementary Figures for "Centimeter-sized Human Vitreous Organoids from Pluripotent Stem Cells"

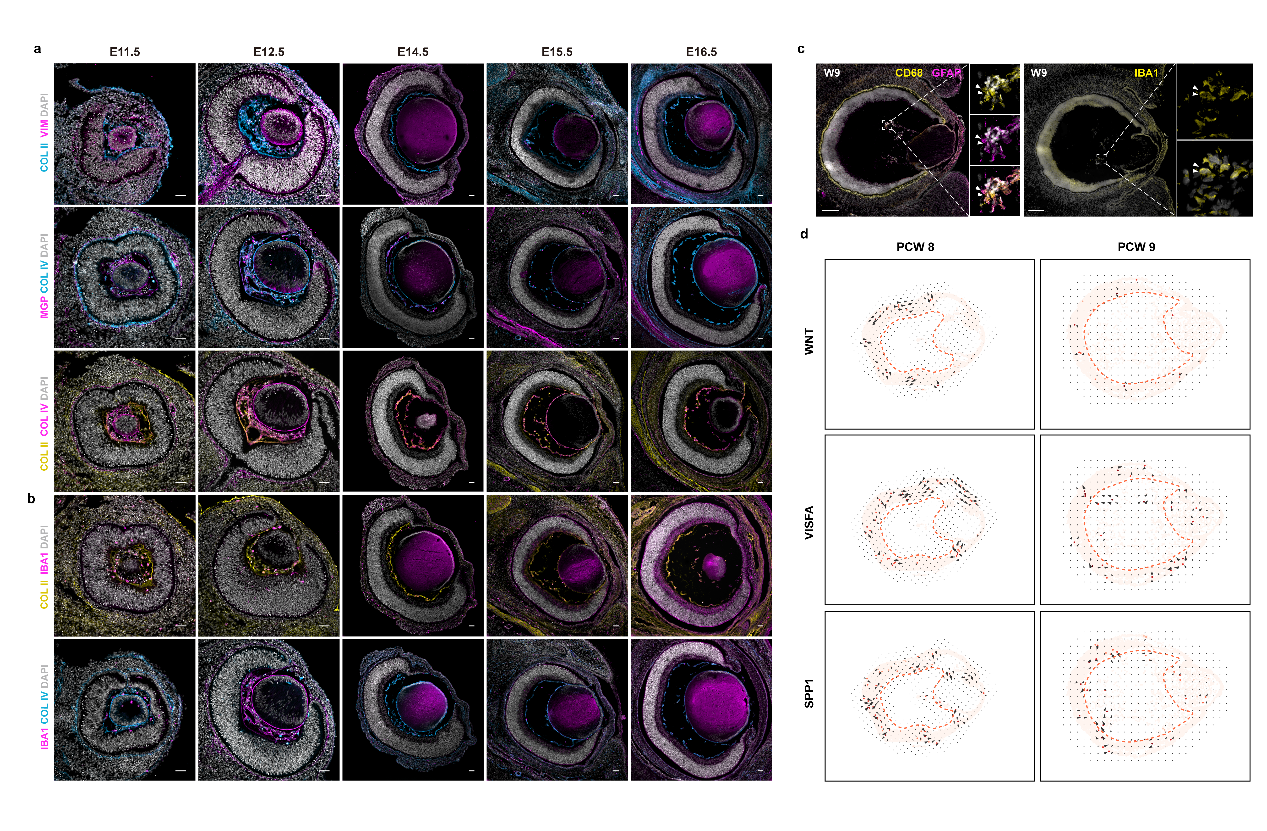

**Extended Data Fig1. Spatiotemporal characterization of vitreous cellular and extracellular matrix dynamics during ocular development. (a)** Immunostaining of murine embryonic eyes (E11.5-E15.5) confirms fibroblast localization (VIM+/MGP+) and extracellular matrix (ECM) deposition (Collagen II/IV) within the developing vitreous. **(b)** Hyalocyte identification (IBA1+/CD68+) in murine embryonic eyes (E11.5-E15.5) reveals temporal coordination with fibroblast activity. **(c)** Human embryonic eye (W9) immunostaining validates conserved hyalocyte markers (IBA1+/CD68+). **(d)** Developmental pathway direction analysis (W8-W9) delineates stage-specific signaling pathway orientations.

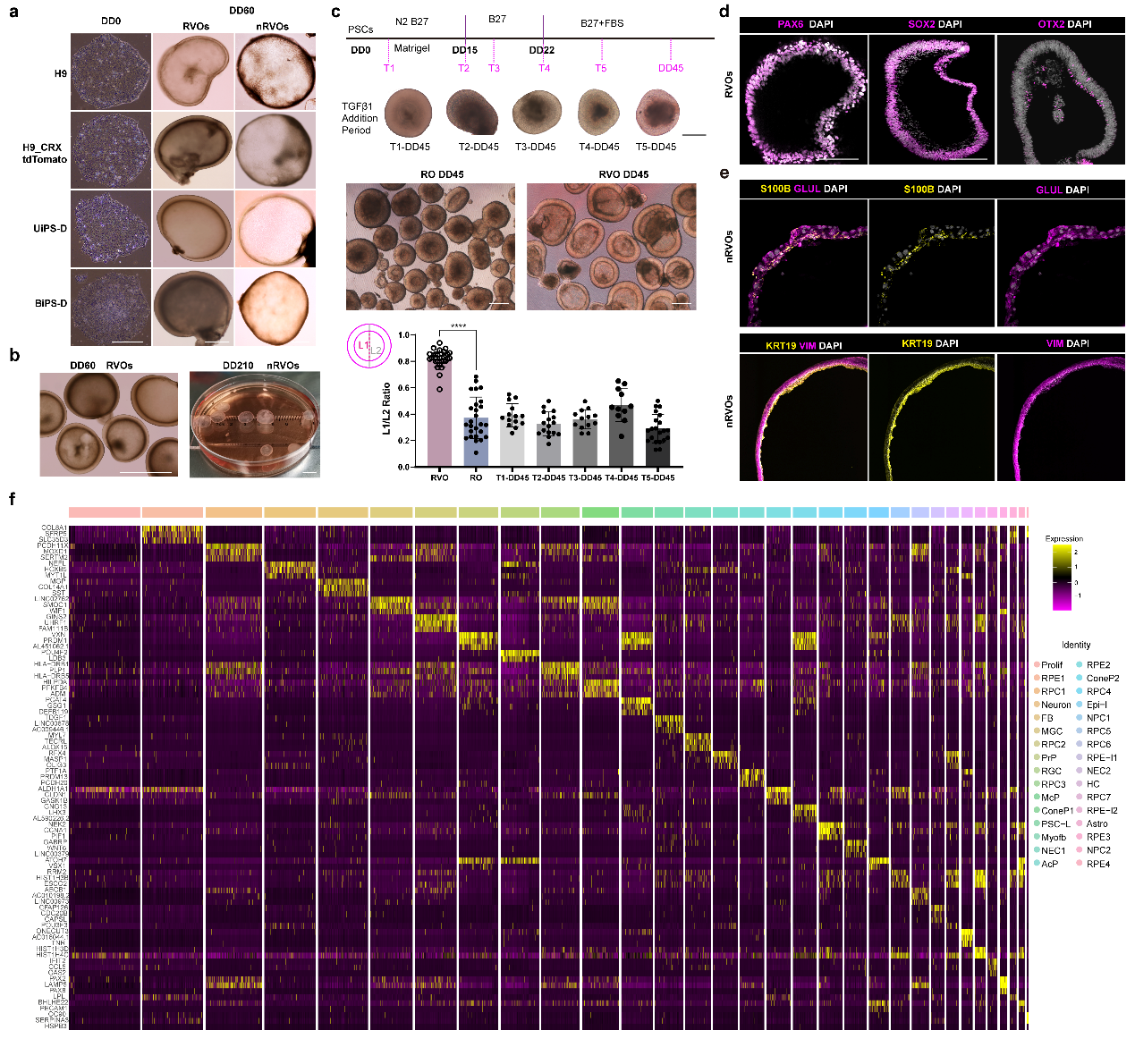

**Extended Data Fig2. Characterization of human pluripotent stem cell-derived vitreous organoids. (a)** Brightfield images of VOs generated from four human pluripotent stem cell lines at differentiation day 60 (DD60). Scale bars, 500 μm. **(b)** Representative brightfield images showing RVOs at DD60 and nRVOs at DD240. Scale bars, 500 μm (RVOs) and 1 cm (nRVOs). **(****c)** TGFβ1 was added to a retinal organoid (RO) protocol at five timepoints (T1-T5) The proportion of organoids containing vitreous is significantly higher in RVO conditions compared to RO conditions, and is unaffected by TGFβ1 in the RO protocol. (**** p < 0.0001).T1-DD45 means TGFβ1 was added from T1 to DD45. The concentration of TGFβ1 is consistent with that in VO differentiation protocol. Scale bars, 200 μm. L1 represents the length of VH and L2 represents length of VO **(d)** Immunofluorescence analysis of cryosectioned RVOs stained with retinal progenitor cell markers PAX6, SOX2, and OTX2. Nuclei were counterstained with DAPI. n=3 biological replicates, Scale bar: 200 μm. **(e)** Immunostaining of nRVO cryosections showing positive expression of and Müller and fibroblast markers S100B, GLUL, KRT19, and VIM. n=3 biological replicates. Scale bars, 50 μm. **(f)** Heatmap displaying the top three marker genes for each identified single-cell cluster.

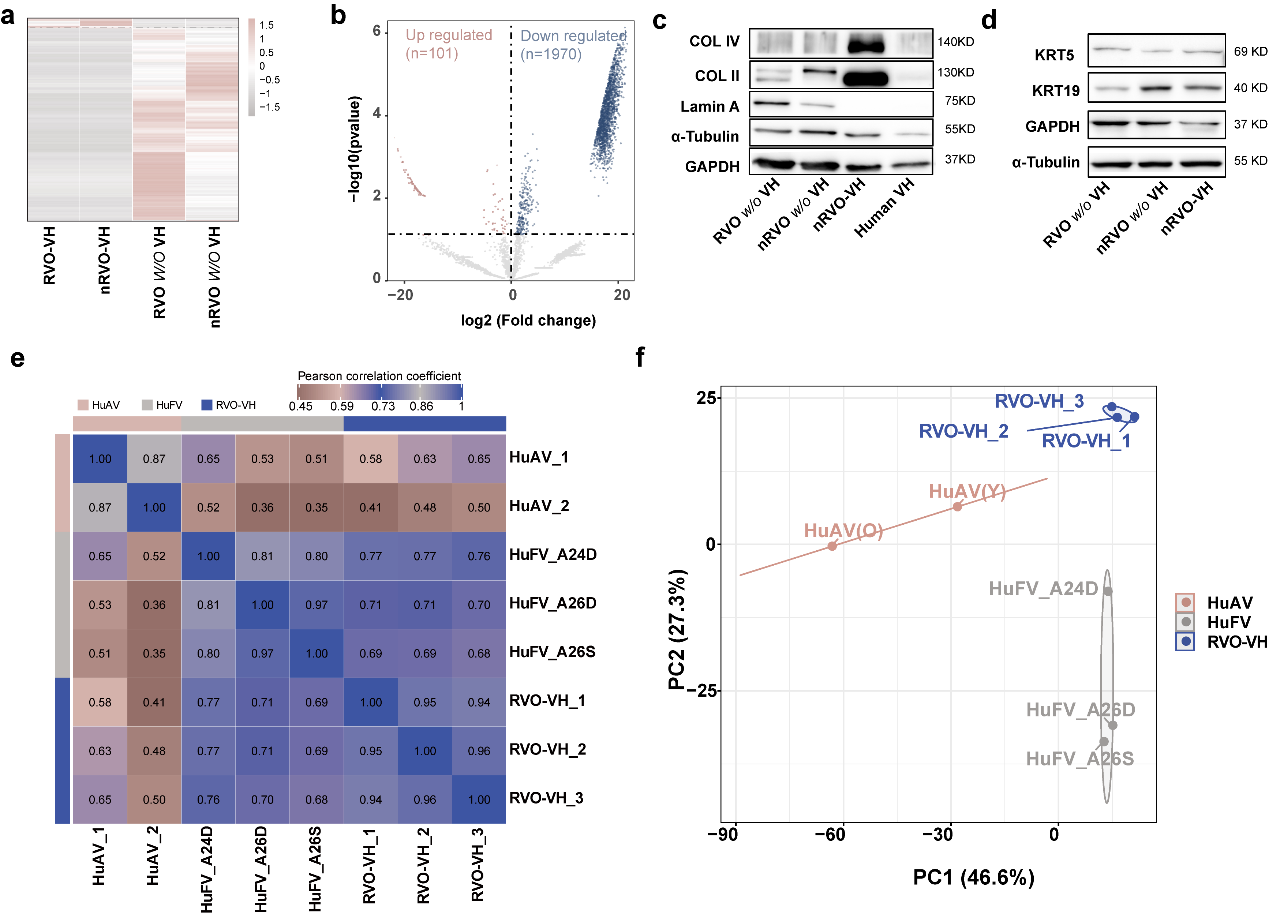

**Extended Data Fig 3. Proteomic profiling and functional validation of VO extracellular matrix components. (a)** Volcano plot showing differentially expressed proteins between vitreous humor (VH) and VO without VH. Dashed lines indicate statistical thresholds (log₂ fold-change >1.5, p < 0.05). **(b)** Gene Set Enrichment Analysis (GSEA) of hallmark pathways enriched in upregulated proteins from the VH group. Normalized enrichment scores (NES) and false discovery rates (FDR) are indicated. **(c-d)** Western blot analysis validating the expression of collagens and structural proteins in nRVO-VH, nRVO w/o VH, and nRVO w/o VH and human VH. Representative blots from two independent experiments are shown. **(e)** Pearson correlation coefficients (r) showing pairwise comparisons of the global proteomic profiles between the indicated VH samples. The high correlation between RVO-VH and fetal VH highlights their molecular similarity. The correlation between left and right eyes from the same adult individual (r=0.97) serves as a technical reference, while the lower inter-individual correlations (e.g., 0.8 between different adults) reflect natural biological variation. **(f)** Principal component analysis (PCA) plot based on the global proteomic profiles of adult, fetal, and RVO-derived VH samples. The close clustering of RVO-VH with fetal VH, and their clear separation from adult VH, demonstrates that the organoid-derived matrix closely resembles the fetal developmental stage.

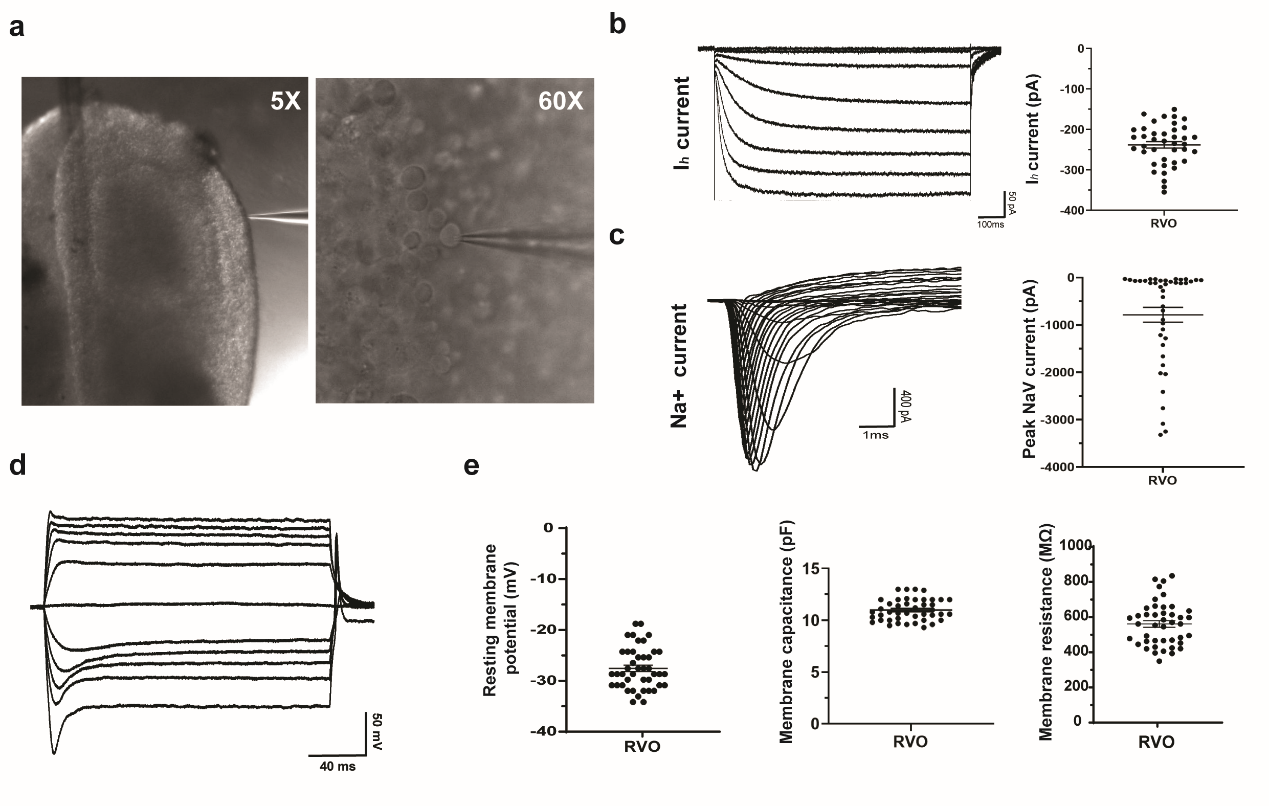

**Extended Data Fig4. Electrophysiological characterization of photoreceptors in DD240 RVOs. (a)** Representative bright-field images of RVOs at 4× magnification showing NR structure and typical morphology of RVO photoreceptors at 60× magnification. **(b)** Hyperpolarization- activated current (HCN current) of photoreceptors and quantifications. Voltage steps from -120 mV to -50 mV in 10-mV increments. Holding potential: -50 Mv. **(c)** Voltage-gated Na^+^ current of photoreceptors and quantification. Voltage steps from -70 mV to +50 mV in 5-mV increments, holding potential: -90 mV. **(d)** Membrane potentials of photoreceptors. Current steps from -100 pA to +100 pA in 20-pA increments. **(e)** Characterization of membrane properties such as resting membrane potential, membrane capacitance, and membrane resistance.

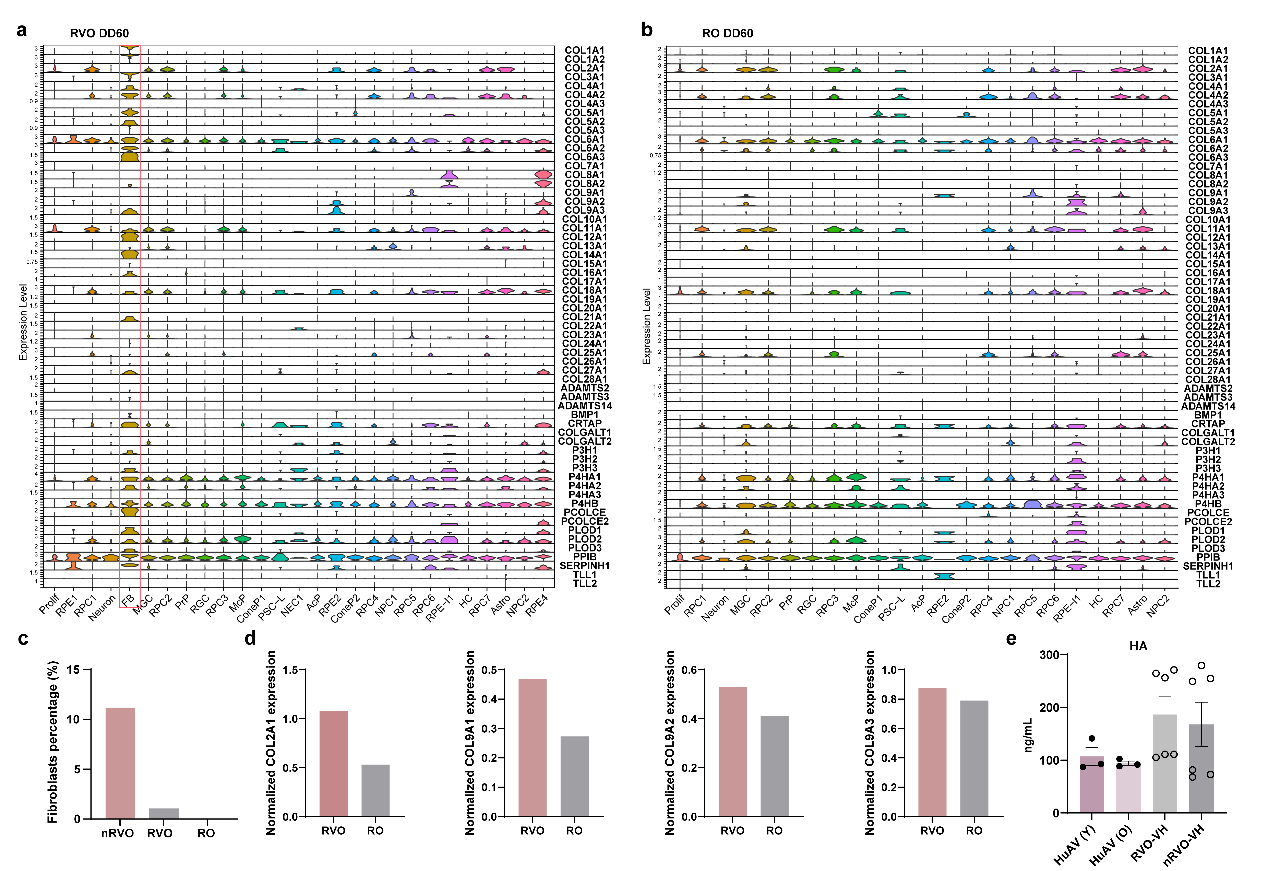

**Extended Data Fig 5. Fibroblasts drive the biosynthesis of the vitreous matrix.** **(a-b)** Heatmap of expression levels for key collagen and collagen-modifying genes in RVOs (a) and ROs (b), underscoring the specific enrichment in fibroblasts.**(c)** ScRNA-seq analysis of fibroblast counts in nRVO, RVO, and RO at day 60. **(d)** Quantitative proteomic analysis of COL2 and COL9 levels in ROs versus RVOs (with VH) (n=2 biological batches per condition). Levels are significantly elevated in RVOs. **(e)** ELISA quantification of hyaluronic acid (HA) concentration in VH from native human adult vitreous (36- and 64-year-old donors) and from in vitro-derived nRVO-VH and RVO-VH (n=2 biological replicates, with n=3 technical replicates each). Data are presented as mean ± SD.
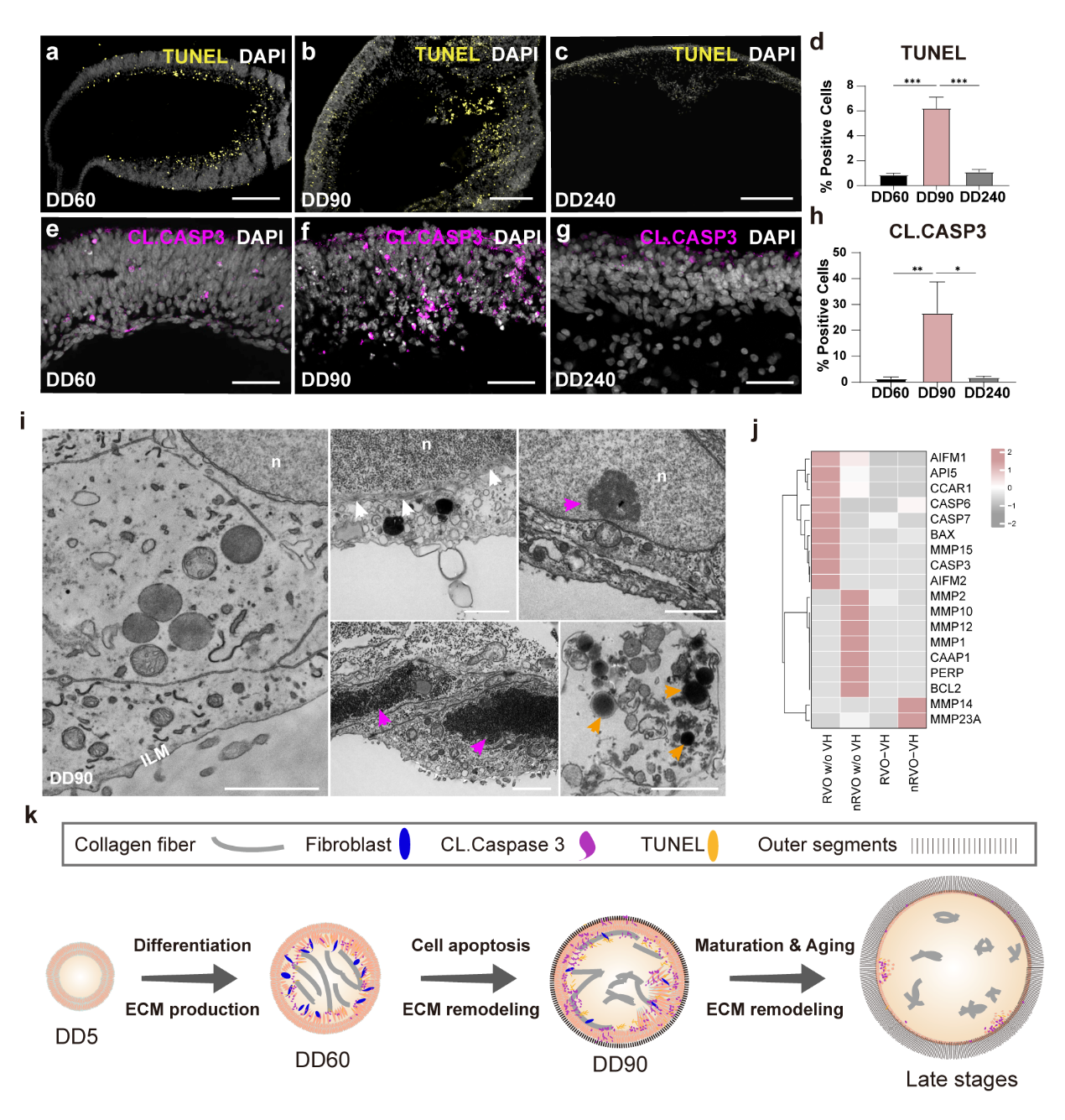

**Extended Data Fig6. Cell apoptosis during vitreous development and remodeling in RVOs.** **(a-c)** TUNEL staining for the apoptotic cells (yellow) in RVOs at different stages. Cells were counterstained with DAPI. Scale bars, 200μm. n=2 biological replicates. **(d)** The statistics results of TUNNEL staining. 10 images of each stage were counted. **(e-g)** Immunostaining of RVOs for cleaved caspase 3 (CL.CASP3), the effector of apoptosis. Cells were counterstained with DAPI. Scale bars, 50μm. n=2 biological replicates. **(h)** The statistics results of CL.CASP3 staining. 10 images of each stage were counted. **(i)** TEM images showing the structure along the ILM and the apoptotic cells in DD90 RVOs. The apoptotic cells shrank (white arrows), with chromatin condensation (red arrows) and apoptotic bodies (orange arrows). n, nuclei. Scale bars, 2μm. **(j)** Heatmap showing the expression of apoptosis-related regulators and ECM modifiers in DD90 VH-depleted RVOs and nRVOs, as well as RVO-VH and nRVO-VH. **(k)** Schematic representation of VH formation during RVO differentiation.

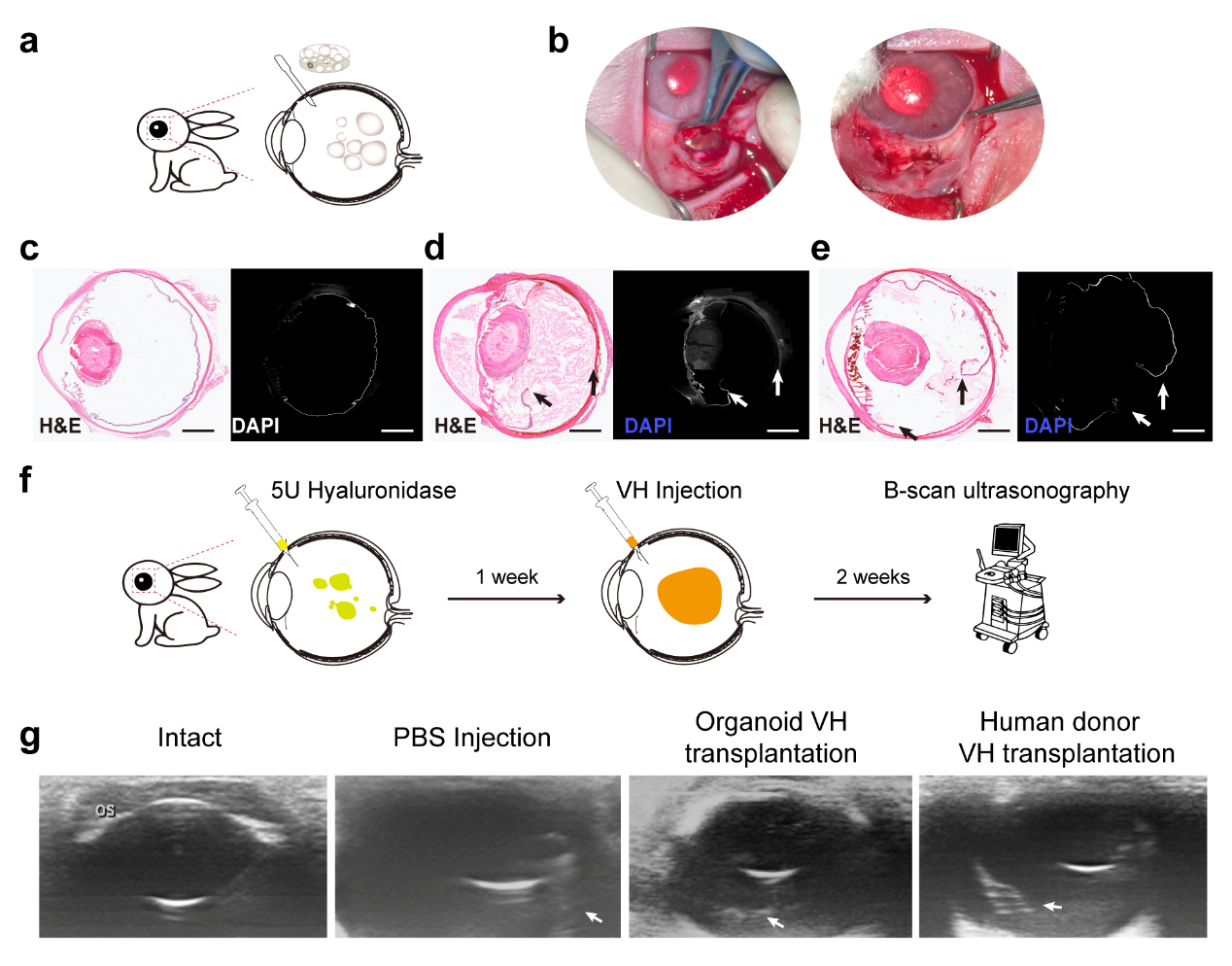

**Extended Data Fig7. Transplantation of VOs, VH from VOs or human donor VH into rabbit vitreous. (a)** Schematic showing the transplantation of RVOs or nRVOs into the left rabbit eyes. **(b)** Images showing the surgical exposure of the vitreous surface and suturing of the wounds after transplantation. **(c)** The H&E and DAPI staining of the rabbit eye without surgery and transplantation. Scale bars, 2.5 mm. **(d)** Transplantation of large volume of nRVOs caused proliferative vitreoretinal diseases (PVD). The large incision surgery procedure caused retinal detachment. The arrows indicated the broken detached retina. Scale bars, 2.5 mm. **(e)** Transplantation of RVOs did not cause PVD, but the large incision surgery procedure caused retinal detachment (as indicated by the arrows). Scale bars, 2.5 mm. **(f)** A schematic shows the transplantation of VH into the vitreous of the left rabbit eyes and the subsequent examination at 2 weeks post-surgery. **(g)** The B-scan ultrasonography for rabbit eyes without surgery and transplantation, transplanted with PBS, VH from VOs, or human donor VH. Arrows indicate the reflection of vitreous floaters.

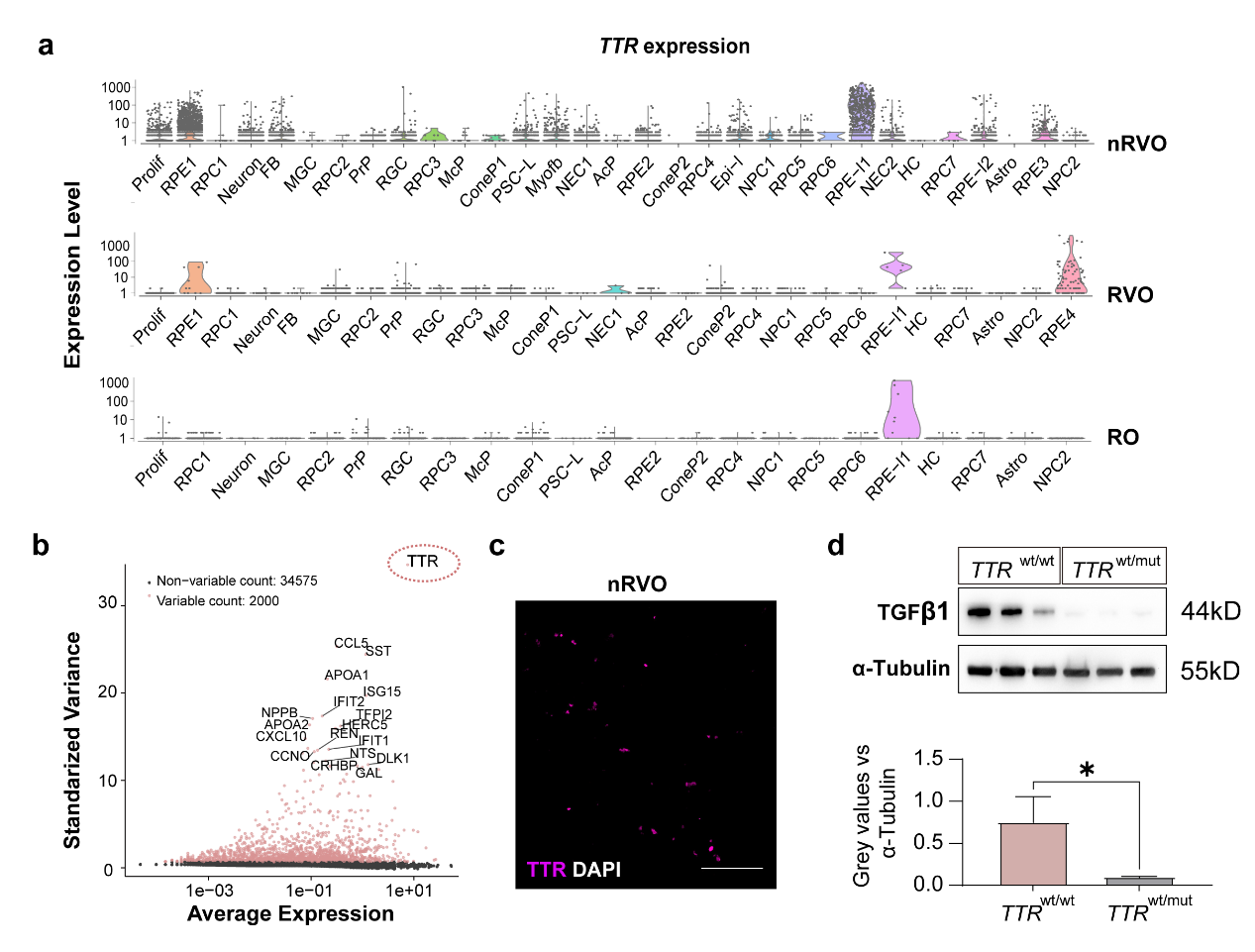

**Extended Data Fig 8. Single-cell transcriptomic profiling and spatial expression of transthyretin (TTR) in vitreous organoids. (a)** TTR expression was predominantly localized to retinal pigment epithelium (RPE)-lineage cells in sc-RNA sequencing results. **(b)** Single-cell RNA sequencing identified TTR as the top contributor gene (by enrichment score) in nRVO-derived vitreous-like organoids. **(c)** Immunofluorescence validation showing diffuse TTR protein distribution across nRVOs, with nuclear counterstaining by DAPI. Scale bars, 100μm. n = 3 biological replicates. **(d)** Western blot analysis of TTR monomer levels in wild-type and TTR-mutant samples, normalized to α-tubulin (loading control). Graph quantifying TTR/α-tubulin ratios, demonstrating significant reduced monomeric TTR levels in mutants compared to wild-type. *, p < 0.05.
