## Supplementary methods for "Centimeter-sized Human Vitreous Organoids from Pluripotent Stem Cells"

**Materials and Methods**

**PSC culture and maintenance**

Three different PCS lines were used in this study. The human blood-derived iPS (BiPS)^41^ and the urine-derived iPS (UiPS)^32^ cell lines were generated in our laboratory. The human ES cell line (H9) was purchased from Wicell research institute. The CRX-tdTomato reporter line (H9-CRX-tdTomato) was genetically modified from H9^42^. These cells were maintained in Vitronectin (Nuwacell Biotechnology, RP01001) in serum-free medium (Nuwacell Biotechnology).

**Derivation of retinal organoids (ROs) and VOs from human PSCs with TGFβ1**

Human ROs were derived from human PSCs based on our previous protocols^43-45^. VOs were generated by the following protocol. Briefly, PSCs were detached by the dispase (StemCell, 07923) and cultured in the stage 1 medium [DMEM/F12 supplemented with 15% Knockout serum replacement (ThermoFisher, 10828028), 2 mM L-Glutmax (ThermoFisher, 35050061), 0.1mM β-mercaptoethanol (ThermoFisher, 21985023), 100ng/μL TGFβ1 (MedChemExpress, HY-P7118) and 10μM Y-27632 (Selleck, S1049)] for 1 week and then replaced with the stage 2 medium [DMEM:F12 (3:1) containing 1% N2 supplement (ThermoFisher, 17502-048), 0.1 mM non-essential amino acids (Sigma, M7145), 2% FBS (Biological industries, 04-002-1A), 100ng/μL TGFβ1] for 3 weeks. After that the cells were refreshed with the stage 3 medium [DMEM:F12 (3:1) containing 2% B-27, 0.1 mM non-essential amino acids, 2 mM L-Glutmax, 0.1mM β-mercaptoethanol, 100mM Taurine, 4% FBS and 200ng/μL TGFβ1] for long-term culture.

**Ethics for medical research and human tissue collection**

Vitreous humor of a healthy donor was collected from Beijing Tongren Hospital. Aborted human embryo samples were collected from West China Airport Hospital. The use of human tissues was approved by the institutional Ethics Committee of Beijing Tongren Hospital, Capital Medical University (TREC2022-KY114 and TREC2024-KY182). Informed written consent was obtained from all participants in accordance with the Declaration of Helsinki guidelines.

**Immunostaining and Western blotting**

VOs were fixed in 4% PFA at 4°C overnight and then embedded in O.C.T (Tissue-Tek) and sectioned. After blocking and permeabilization, sections were incubated with primary antibodies overnight. Antibodies used were listed in detail in the supplementary material (Supplementary Table 1). Images were acquired using a confocal microscope (Olympus). For immunoblotting, clinical vitreous samples were concentrated (ThermoFisher) and VOs were homogenized in the RIPA buffer at 4°C before boiled with loading dye (ThermoFisher) and 2-Mercaptoethanol (ThermoFisher). Then samples were loaded into an SDS-PAGE gel. After electrophoresis and immunoblotting, protein bands on membranes were imaged using a Li-COR Odyssey Infrared Imaging System.

**Transmission electron microscopy analysis**

The DD90 or DD240 VOs were fixed with 4% paraformaldehyde, 2% glutaraldehyde in 0.1 M cacodylate buffer (pH 7.4) at 4°C overnight and then postfixed in 1% osmium tetroxide. Samples were dehydrated in a gradient series of ethanol and acetone, followed by resin infiltration. Ultrathin (100 nm) sections were imaged with a Zeiss electron microscope.

**Scan electron microscopy analysis**

The DD240 RVOs were fixed with 2.5% glutaraldehyde in sodium cacodylate buffer at 4°C overnight. The samples were then dehydrated through a gradient series of ethanol and transferred into a Leica EM CPD300 critical-point dryer. After critical-point drying, the samples were mounted on aluminum stubs and sputter coated with a thin layer of gold. The samples were viewed on a Hitachi-SU8010 field emission scanning electron microscope at an accelerating voltage of 5 kV.

**X-ray micro-CT scanning**

VOs were pretreated with 2% phosphotungstic acid for 1 hour at room temperature and then dehydrated through a gradient series of ethanol and kept in the absolute ethanol. VOs were loaded in 200 μL pipet tips in absolute ethanol, sealed with paraffin oil and parafilm and attached on tips. The X-ray source was operated at a voltage of 60 kV and a power of 5 W during the scanning process. The CCD optical objective (4X) was used for high resolution images acquisition. The scanning resolution was set to 1.0 μm per vortex under a binding mode of 1.0 × 1.0 for DD60 organoids, 3.0 μm per vortex under a binding mode of 3.0 × 3.0 for DD180 organoids, and 0.6 μm per vortex under a binding mode of 0.6 × 0.6 for zoom in parts. The 3D reconstruction of obtained images and segmentation for each specimen was performed using Dragonfly software (version 2021).

**Second-harmonic generation (SHG) imaging**

The DD60 RVOs were chosen for SHG imaging using two-photo microscope. Before imaging, organoids were fixed with 4% paraformaldehyde and then washed three times with DPBS. The RVOs were imaged using a Zeiss LSM 980 confocal microscope that equipped with a Ti:Sa Chameleon multiphoton tunable laser (Coherent, Santa Clara, CA, USA) at 800 nm and a 20× water immersion objective. The resulting images were processed using Zen software (Zeiss Microscopy, Jena, Germany).

**RNA-seq analysis**

VOs were homogenized in TRIzol reagent and total RNA was extracted. RNA integrity and concentration were assessed using an RNA Nano 6000 Assay Kit and a Bioanalyzer 2100 system (Agilent Technologies). A total amount of 2 μg RNA was used for preparation of RNA sequencing libraries with NEBNext Ultra RNA Library Prep Kit (NEB, E7530L), following the manufacturer’s recommendations. All RNA-seq samples were sequenced by Illumina Nova 6000 with pair-end 150-bp read length. Quality control of the reads generated from RNA-Seq was performed using fastqc^46^. All reads were mapped to human genome version hg38 by hisat2 (version 7.5.0) with default settings ^47^. Read counts were calculated using feature Counts^48^. Differential gene expression was determined with edge R^49^ with a cutline of log2 (Experimental group/Control group) >1 and p < 0.05. Data visualization was carried out with R language and packages.

**Label-free quantitative proteomic analysis**

VOs were homogenized in the RIPA buffer at 4°C and the supernatants were collected after centrifuging. Samples in liquid format were digested by trypsin, and then identified and quantified by nanoLC-MS/MS platform. The protein modifications were carbamidomethylation (C) (fixed) and oxidation (M) (variable); the maximum missed cleavages were set to 2; the precursor ion mass tolerance was set to 20 ppm, and MS/MS tolerance was 0.6 Da. Only high confident identified peptides were chosen for downstream protein identification analysis. Protein database was based on the species of the samples using Maxquant (1.5.6.5).

**Patch-clamp recordings**

Three RVOs were collected for patch-clamp recording. RVOs were transferred to the patch-clamp recording chamber (5% CO2, 95% O2) filled with bubbled Ames' solution containing 120 mM NaCl, 22.6 mM NaHCO3, 6 mM D-glucose, 3.1 mM KCl, 1.2 mM MgSO4, 1.1 mM CaCl2, 0.5 mM KHPO4, and 0.5 mM L-glutamine. Recording pipettes were made using a Sutter P-1000 puller (Sutter Instruments) to pull glass capillaries (Sutter Instruments, California: OD 1.5 mm, ID: 0.86 mm) to a tip resistance of 3-8 MΩ when filled with the internal solution containing 110 mM KCl, 13 mM NaCl, 2 mM MgCl2, 1 mM CaCl2, 10 mM EGTA, 10 mM HEPES, 5 mM ATP-Na, and 0.5 mM GTP-Na, with the pH adjusted to 7.2 with KOH. Recordings were obtained using an AxoPatch 200B amplifier and a Digidata 1550B digitizer (Molecular Devices, California). Data acquisition was performed using Axon Clampex software (Molecular Devices, v10.6), with signals low-pass filtered at 2 kHz and sampled at 10 kHz.

**Transplantation of vitreous organoids or vitreous humor into rabbit eyes**

Four rabbits (weighing 2.0–3.0 kg; Beijing HealthOlight Technology Co., Ltd, China) were used in this study. All procedures were performed on the right eyes and the left eyes were used as the control. Topical tropicamide was applied three times with an interval of 5 minutes to dilate the pupil. For transplantation of vitreous organoids, the rabbit vitreous was partially removed by surgery under anesthesia, and an equal volume of vitreous organoids were transplanted. For transplantation of vitreous humor, a vitreous liquefaction model was created by injecting 5U (100 μL) hyaluronidase into the central rabbit vitreous. After 1 week, one eye was used as a blank control, and part of the vitreous humor in the other eye was replaced with 100μL PBS, vitreous humor from organoids, or donated healthy human vitreous humor. Two weeks later, the rabbit vitreous was examined by B-scan ultrasonography.

**Generation of hESCs carrying a *TTR* mutation**

The H9 ES cells (from ATCC) were cultured in a 6-well plate for 2 days and then transfected with Lipofectamine Stem (ThermoFisher, STEM00008) when they reached approximately 30% confluence. A total of 600 ng of Cas9 plasmid and 300 ng of pegRNA-expressing plasmid (guide RNA sequence: tgcacggccacattgatggc) was transfected into cells according to the manufacturer’s recommendations. Two days after transfection, 1μg/ml puromycin (InvivoGen, ant-pr-1) was added to the medium. Cells were incubated for another 4 days and then collected for high-throughput sequencing. After the *TTR* mutation was identified, single clones were selected, identified, and expanded.

**Stereo-seq library preparation and data processing**

We employed the STOmics Gene Expression kit S1 (BGI, 1000028493) following the manufacturer’s protocol V1 ^50^. Initially, tissue sections were mounted onto the Stereo-seq chip and incubated at 37°C for 3 minutes. The sections were then fixed in methanol at -20°C for 40 minutes before being stained with a nucleic acid dye (Thermo Fisher, Q10212) and imaged using a microscope under FITC illumination. Subsequent to imaging, the sections were washed with 0.1× SSC buffer (Thermo, AM9770) and subjected to permeabilization using 0.1% pepsin in a 0.01 M HCl solution for 6 minutes at 37°C. The RNA released from the tissues was reverse transcribed using SuperScript II (Invitrogen), and the sections were rinsed twice with 0.1× SSC buffer. Following tissue digestion with a removal buffer, the chips containing the synthesized cDNA were incubated with Exonuclease I at 55°C overnight, followed by a wash with 0.1× SSC buffer. The resulting cDNA underwent amplification and fragmentation by Tn5 transposase, with a subsequent re-amplification step using KAPA HiFi Hotstart Ready Mix. Finally, the amplified products were purified and processed for DNA nanoball (DNB) generation, after which sequencing was carried out on the MGI DNBSEQ-T1 platform.

For Stereo-seq analysis, the raw fastq files were structured such that read 1 comprised the CID (bases 1–25) and MID (bases 26–35), while read 2 contained the cDNA sequences. The initial step involved aligning these raw reads to a predetermined coordinate matrix of the in situ captured chip, permitting a single-base mismatch. Reads with UMIs that either included ambiguous 'N' bases or had quality scores below 10 were discarded. The remaining high-quality reads were then mapped to the mm10 genome reference using STAR2 ^51^, and only those with a MAPQ score of at least 10 were kept for further analysis. Next, the mapped reads were quantified and assigned to corresponding genes with handleBam, resulting in a gene expression profile matrix. This workflow was executed using the publicly available SAW pipeline (<https://github.com/STOmics/SAW>). Finally, the gene expression matrix was normalized via SCTransform, and unsupervised clustering was performed using Seurat to classify distinct cell types within non-overlapping cellbins, where transcripts from the same gene were aggregated for subsequent analysis.
